# Evolution of cullin E3 ubiquitin ligases and function in trypanosomes

**DOI:** 10.1101/2023.07.24.550360

**Authors:** Ricardo Canavate del Pino, Martin Zoltner, Kayo Yamada, Erin R. Butterfield, Mark C. Field

**Affiliations:** School of Life Sciences, University of Dundee, Dundee, UK; Charles University in Prague, Faculty of Science, Department of Parasitology, Vestec 252 42, Czechia; Institute of Parasitology, Biology Centre, Czech Academy of Sciences, České Budějovice, Czechia

**Keywords:** cullin, ubiquitin, E3 ligase, protein turnover, evolution, affinity isolation, cryomilling, eflornithin, ornithine decarboxylase, trypanosoma, proteomics

## Abstract

Post-translational modifications (PTMs) modulate protein function, with ubiquitylation a pre-eminent example with major roles in protein turnover. Ubiquitylation utilises a ligase enzyme cascade for conjugation of ubiquitin to client proteins and cullin-RING ligases are amongst the most complex known. We reconstructed evolution of cullin-RING E3 ubiquitin ligases across eukaryotes and experimentally characterised two cullin complexes in trypanosomatids, a taxon highly divergent from animals and fungi. We find considerable diversity within cullins and, in particular, trypanosomatids share only a minority of cullins with other lineages. Furthermore, we identify expansions in cullin client adaptor protein families, novel client adaptors and demonstrate client specificity. Finally we show that ornithine decarboxylase (TbODC), an important target of the drug trypanosome eflornithine, is a substrate for TbCul-A and overturn earlier models for eflornithine specificity. These studies highlight lineage-specific roles for cullin E3s and their contributions towards eukaryotic complexity.

## Introduction

Maintaining a steady state, and facilitating rapid changes in the proteome are essential for cellular viability. Of many mechanisms involved the attachment of ubiquitin to client proteins is most prominent. Ubiquitylation has multiple roles, with decreased stability *via* proteasome targeting and/or altered sub-cellular localisation as major functions (Komander and Rape 2012). Initially, free ubiquitin is activated by an E1 ubiquitin transferase, followed by transfer *via* an E2 conjugating enzyme to an E3 ligase. The final step involves recognition by the E3 ligase of a client protein and covalent attachment of ubiquitin *via* the C-terminal carboxyl to NH_2_ lysine side chains, creating an isopeptide bond (Buetow and Huang 2016).

Most E3 ligases are classified as RING, HECT, U-box, PHD-finger or RING between RING (RBR), based on domain architecture. The *Homo sapiens* genome encodes over 600 RING, 30 HECT and 12 RBR proteins (Morreale and Walden 2016), and amongst the RING class, ligases are notably diverse, characterised by a zinc finger RING domain and recruitment into multisubunit complexes. The *H. sapiens* SCF (<u>S</u>kp/<u>c</u>ullin/<u>F</u>-box) is the prototypical cullin-RING complex and possesses a core cullin scaffold protein binding RBX1, a RING E3 ligase at one end and SKP1/F-box client protein adaptors at the opposite end (Zheng et al., 2002). A repertoire of client adaptor subunits allows cullin ligases to recruit and recognise numerous clients, and activity is tightly regulated *via* neddylation and phosphorylation (Baek et al. 2020, Clague et al., 2015). The basis of specificity, origins and adaptations of ubiquitylation pathways remain poorly characterised. Significantly, the anaphase promoting complex (APC) and replisome also contain a cullin/ Rbx ubiquitin ligase at their core and govern turnover of critical proteins during mitosis (Alfieri et al., 2017, Jenkyn-Bedford et al., 2021).

Originally considered eukaryotic-specific, ubiquitylation predates eukaryogenesis (Grau-Bové et al., 2015, Fuchs, et al., 2018, Barlow et al., 2023). Identification and biochemical characterisation of a complete ubiquitylation pathway in Archaea is consistent with the near universal presence of the proteasome. Moreover, ubiquitylation systems are present in many bacterial lineages, most likely resulting from lateral gene transfer (Levin-Kravets et al., 2018, Barlow et al., 2023). The ubiquitylation system has considerably expanded and diversified in eukaryotes, with a repertoire of over 600, 500 and 100 E3 ligases in the *H. sapiens, Arabidopsis thaliana* and *Saccharomyces cerevisiae* respectively. Ubiquitylation is present in all major eukaryote lineages, but systematic analysis has focused on animals, higher plants and fungi (Morreale and Walden 2016, Jiménez-López et al., 2018, Finley et al., 2012). *In silico* comparative studies have been quite limited, in part due to the complexity of the ubiquitylation machinery, but some patterns are recognised. Specifically, expansions in RING E3 ligases are common, while HECT and RBR families are generally smaller; the latter of similar size as the E1 and E2 enzyme families, with typically two to 40 paralogs respectively in most organisms (Stewart et al., 2016, Clague et al., 2015). However, the presence of lineage-specific components or distinct configurations in terms of subunit connectivity or components within otherwise conserved ubiquitylation pathways remains largely unaddressed.

Trypanosomes, members of the Kinetoplastida within the Discoba lineage, are a model system for evolutionary cell biology, combining tractability, relevant phylogenetic position, divergent biology and comparative simplicity in genome and proteome size. Discoba likely represent an early branch following eukaryogenesis and hence valuable for illuminating processes following eukaryogenesis (Burki et al., 2020). Unsurprisingly, ubiquitylation in trypanosomes regulates the cell cycle and is essential for cellular homeostasis (Rojas et al., 2017, Hu et al., 2017, Benz and Clayton 2007, Damianou et al., 2020, Burge et al., 2022). Both the proteasome and ubiquitylation are involved in targeting and turnover of trypanosome surface proteins (Listovsky et al., 2011, Sharma et al., 2015, Chung, et al., 2008, Quintana et al., 2020), and a canonical ESCRT system functioning in late endocytic trafficking is present (Leung et al., 2008, Silverman et al., 2013, Venkatesh et al., 2018). Further, multiple genetic screens implicate ubiquitylation in stress responses (including therapeutic agents) and gene expression (Alsford et al., 2009). For example, key elements including E3 ligases are drug resistance-associated genes, with modes of action remaining to be explored (Currier et al., 2018). Some trypanosomatids secrete and direct ubiquitylation-regulating proteins into the host cell nucleus, although precise functions remain unknown (Hashimoto et al., 2010). Previous work has addressed the potential functions of two trypanosome cullins (referred to here as TbCul-A and TbCul-E), although neither the composition of these complexes nor their impact on the proteome was investigated (Rojas et al., 2017, Hu et al., 2017).

As the evolutionary origins of E3 ligase complexes are difficult to reconstruct, we considered trypanosomes to examine diversity and function across eukaryotes. To address the diversity and evolutionary history of cullin complexes we used phylogenetic reconstructions to generate an in-depth view of cullin evolution. By high efficiency, single step immunopurification we also characterised trypanosome cullin complex composition and identified lineage-specific adaptations, while silencing allowed examination of the functions of selected cullins. We find considerable under-appreciated complexity within cullin ligases, suggesting considerable inter-taxon variation and identify a specific role for a cullin complex in trypanosome drug sensitivity.

## Results

### Trypanosomes have seven cullin paralogs, the majority lineage-specific

The cullin-RING family stands out due to the large cohort of proteins deployed by the system for recruitment, recognition and transfer of ubiquitin to client proteins. An array of adaptor subunits recruit specific client proteins; both the RING E3 ligase and these adaptor subunits are physically connected by the cullin (**Figure 1**). The number of cullins encoded in the genome varies between species, with eight in *H. sapiens* and five in *A. thaliana*, each forming distinct complexes with specific families of substrate adaptors (Shabek and Zheng 2014, Sarikas et al., 2011).

**Figure 1.**
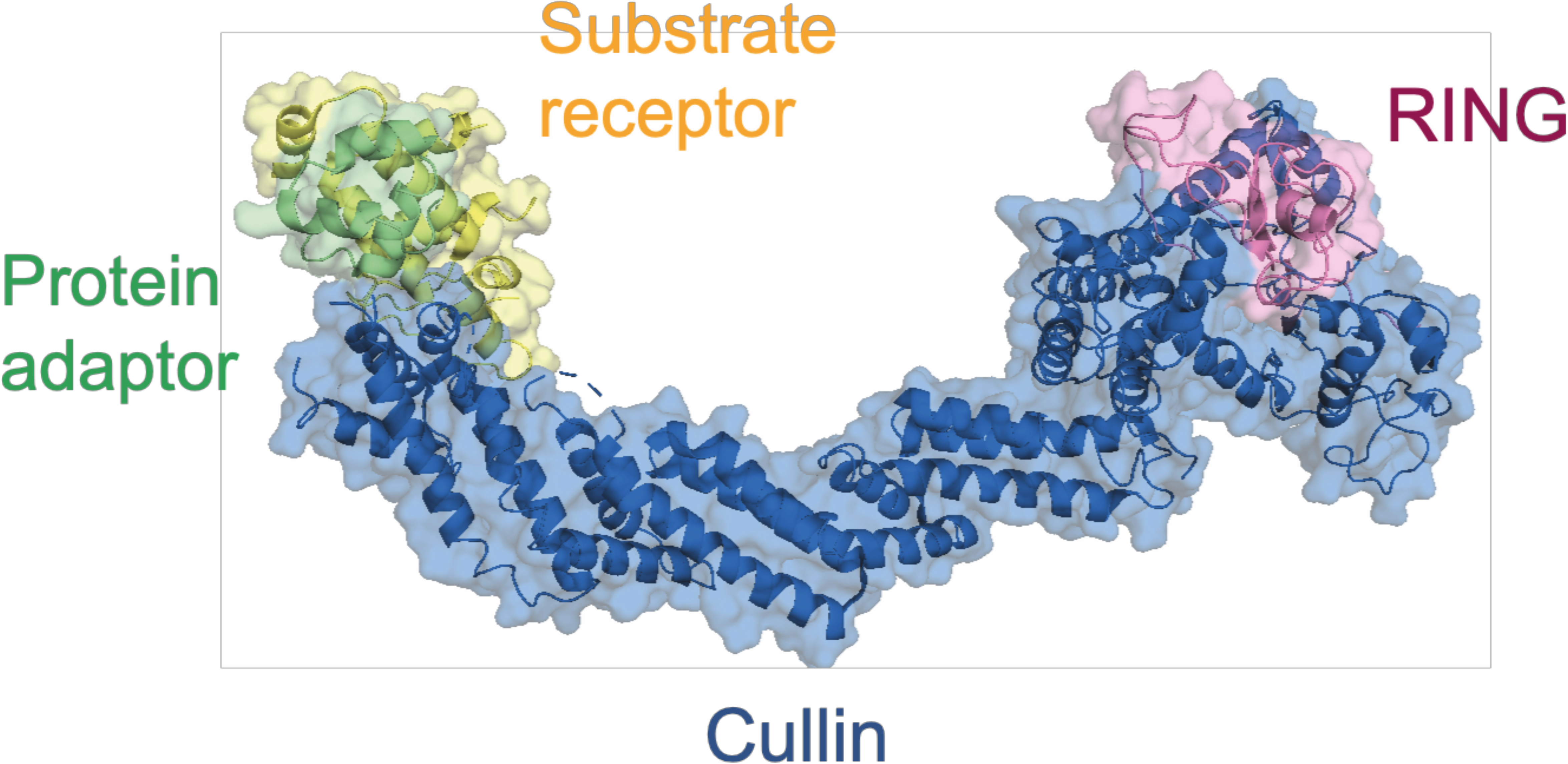
The Cullin-RING complex. The cullin protein (blue) acts as scaffold protein to bridge the E3 ligase RING (pink) to the protein adaptor (green) and substrate receptors (yellow), which recruit and provide selectivity for the ubiquitylation substrates (PDB: 1ldk).

Phylogenetic reconstruction of cullin evolution based around animals, fungi and plants identified three ancestral clades Culα, Culβ and Culγ (Sarikas et al., 2011). Modern cullin ancestry is reflected in substrate receptors associated with each complex; hence descendants from Culα interact with F-box proteins, Culβ with DCAF domain-containing subunits and Culγ with BTB proteins. Significantly, kinetoplastids are reported to encode five, six or seven cullin genes, but evolutionary origins are unclear due to sequence divergence (Marín 2009, Rojas et al., 2017), hampering a pan-eukaryotic phylogeny.

For systemic analysis of the cullin E3 ligase family we used ScrollSaw, with particular emphasis on kinetoplastids (Elias et al., 2012). Our analysis spanned over 130 species and ∼900 protein entries, including the anaphase promoting complex (APC) subunit 2, a highly conserved cullin-RING E3 ligase, as outgroup. Phylogenies were generated for each eukaryotic supergroup and sequences with shortest branches selected for pan-eukaryotic reconstruction. We provisionally named the trypanosome sequences as TbCul-A, TbCul-B etcetera to avoid assumptions of orthology with other organisms. The resulting phylogeny provides a broad reconstruction of the cullin family across eukaryotes, confirms the ancient origin of three ancestral clades and clarified contradicting reports on the number of cullins in trypanosomes, for which it is clear that seven are present and consistent with Rojas (Rojas et al., 2017) (**Figure 2**).

**Figure 2.**
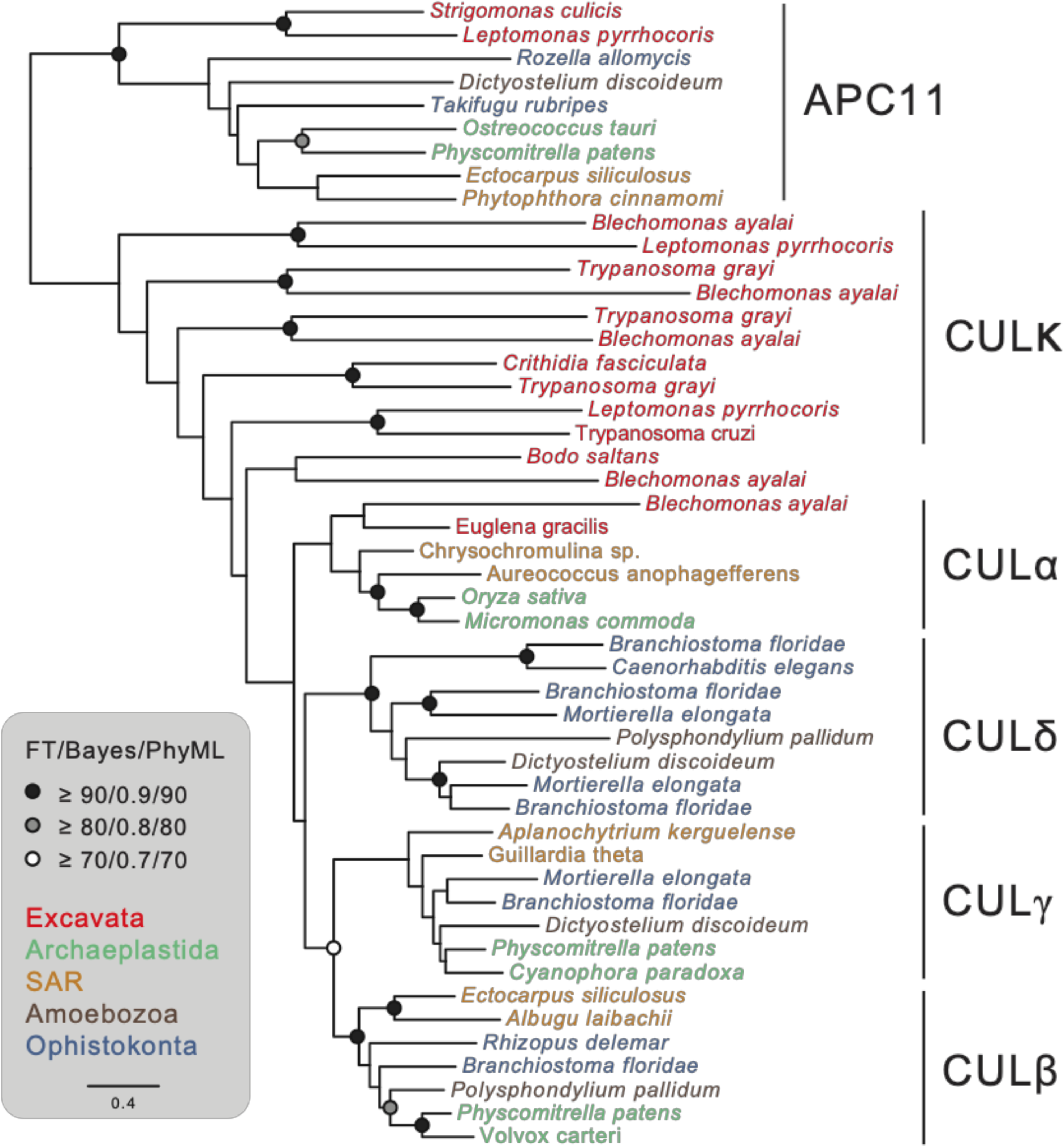
Four ancient clades and the great expansion of cullins in Excavata. Modern cullins in Archaeplastida and Opisthokonta have been proposed to originate from three ancient genes, culα, culβ and culɣ. Each cullin clade has distinct substrate receptors and appear to have expanded independently within the eukaryotic lineages. Although cullins are present in the remaining eukaryotic supergroups, their number and origins are unknown. Reconstructing the phylogenetic tree of cullins with representation from every eukaryotic supergroup reflects the proposed notion of three ancient clades (CULα, CULβ and CULγ). An additional amorphea clade was identified (Culδ) which likely arose post-LECA. Only one of the seven cullins from Excavata, Cul-A, forms part of the major clades and the remaining, appear to be kinetoplastid specific as CULκ (κ for kinetoplastida). Taxa are colour-coded based on supergroup, indicated on the bottom right. The Anaphase Promoting Complex subunit 2 (APC2) was used as an outgroup to root the tree. The support of the nodes, a measurement of confidence, is shown when >70% in each of the three inference methods of phylogeny. The scale bar at the bottom indicates the amino acid substitution rates.

Besides the APC2 outgroup, the pan-eukaryotic phylogeny possesses four major clades, formed by clusters containing CUL-3 and CUL-4 orthologs, consistent with earlier analysis and designation as clades Culα, Culβ and Culγ (Marín 2009). However, CUL-1 and multiple Cul-1-derived cullins from Opisthokonta did not cluster with Culα and instead formed a fourth clade, which we termed Culδ. All cullins from the Amoebozoa and SAR are grouped into one of the three ancestral clades (α, β, γ) indicating that the ancestral clades span eukaryotic supergroups beyond Archaeplastida and Opisthokonta and hence are more ancient than previously suggested (Marín 2009). The branching order of these clades suggests that Culα likely diverged first, followed by Culβ and Culγ, while Culδ is taxon-restricted and may have arisen post-LECA.

Of the seven cullins identified in Discoba, Cul-A clustered within the Culα clade. In contrast, the remainder, Cul-B to Cul-G, were excluded from Culα, Culβ, Culγ or Culδ, indicating they are either lineage-specific or so diverged they cannot be placed with confidence within pan-eukaryotic clades. Long branch lengths suggest phylogenetic artefact, but additional analyses, by removing subsets of sequences (not shown), support this conclusion (but see below). We chose to designate this cluster as Culκ, and suggest that many cullins in trypanosoma, and possibly other Discoba, have lineage-specific origins, and hence possible functional divergence, *albeit* that we cannot fully exclude divergence preventing assignment to a pan-Eukaryotic clade. We suggest to name trypanosoma cullins with letters as nomenclature as using numbers implies orthology with pan-eukaryotic cullins, which is unsupported.

### E3 complex composition in trypanosomes reveals lineage-specific and conserved cullins

Based on current paradigms each of the three primordial cullins recruits one family of client recognition adaptors. The Culδ clade possess novel receptors including VHL and SOC proteins (Marín 2009; Sarikas et al., 2011). Only one cullin from *T. brucei* has been studied in detail to date and this complex possesses substrate adaptors orthologous to DDB1 and DCAF, and was designated as TbCul-4 (Hu et al., 2017). Members of the Culα clade include F-box proteins and Skp1, but the remaining trypanosome cullins are completely uncharacterised.

To address this we used immunoisolation/mass spectrometry to determine the composition of each cullin complex. Six out of seven cullins (TbCul-A, TbCul-C, TbCul-D TbCul-E, TbCul-F and TbCul-G) were successfully endogenously tagged at the C-terminus with a 3×HA-epitope tag and confirmed by Western blotting (**Figure S1**), with TbCul-C and TbCul-F present at significantly lower abundance. Attempts to tag TbCul-B were unsuccessful, and also refractory by high throughput efforts (Billington et al., 2023, tryptag.db). Given the dynamic nature of client adaptor/cullin interactions we used cryo-milling for rapid disruption of cells under near native conditions (Obado et al., 2018). Multiple buffer conditions were evaluated for each isolation and captured proteins visualised using silver-stained SDS-PAGE, allowing identification of appropriate biochemical conditions to obtain an optimal signal to noise ratio **(Figure 3).** Immunocomplexes were analysed using LC-MS/MS and compared with an immunoisolation from untagged parental cells under identical conditions. All analyses were performed in at least triplicate and analysed using MaxQuant and Perseus (Zoltner et al., 2020).

**Figure 3:**
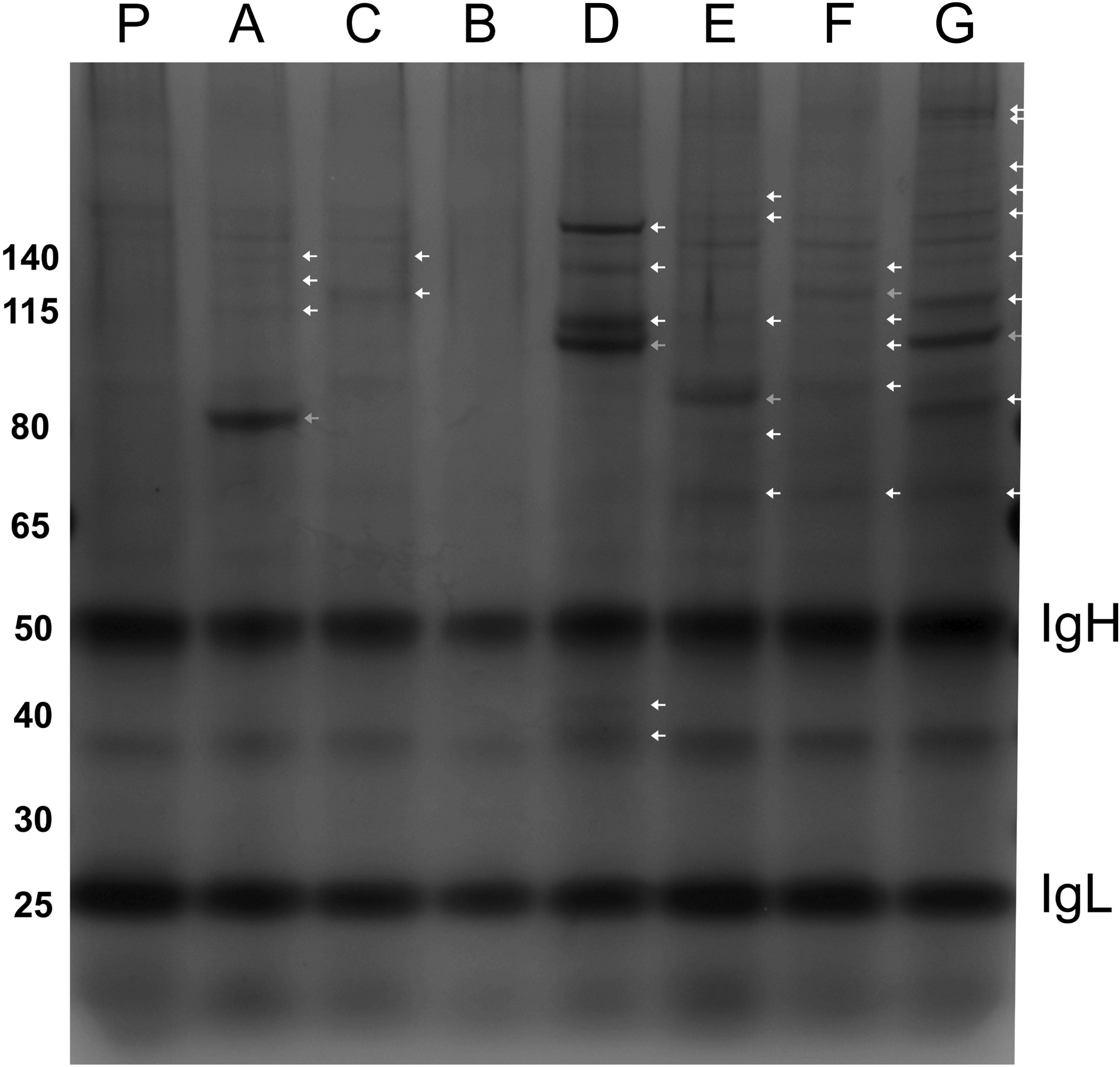
Interacting partners of the *Trypanosoma brucei* cullins reveal the evolution of the complexes. Silver stained SDS-PAGE gel of the elute from the affinity purified 3×HA-cullins from *T. brucei* alongside their interactors. Each band specific to the tagged cell line (i.e. not found in the control) has been labelled based on the predicted molecular mass of the interactors identified through mass spectrometry (Figure 4). Subsequently, the cullins were categorised as public when its interactors indicated they form complexes conserved among eukaryotes or as private cullins, when they appear exclusive to the Kinetoplastida.

We identified multiple F-box and DCAF paralogs amongst proteins enriched in the TbCul-A and TbCul-D isolates respectively **(Figure 4)**, together with orthologs of Skp1, DDB1 and Rbx1 in all cullin isolations (Table 1). Interacting partners of TbCul-D are consistent with previous reports (Hu et al., 2017), while TbCul-A interaction partners, specifically F-box proteins, confirms inclusion within the Culα clade. The presence of DCAF adaptors associated with TbCul-D is consistent with this complex being equivalent to TbCul-4. By contrast, apart from Skp1 paralogs, remaining analyses failed to identify canonical substrate recognition subunits defining Culα, Culβ or Culγ clades.

**Figure 4:**
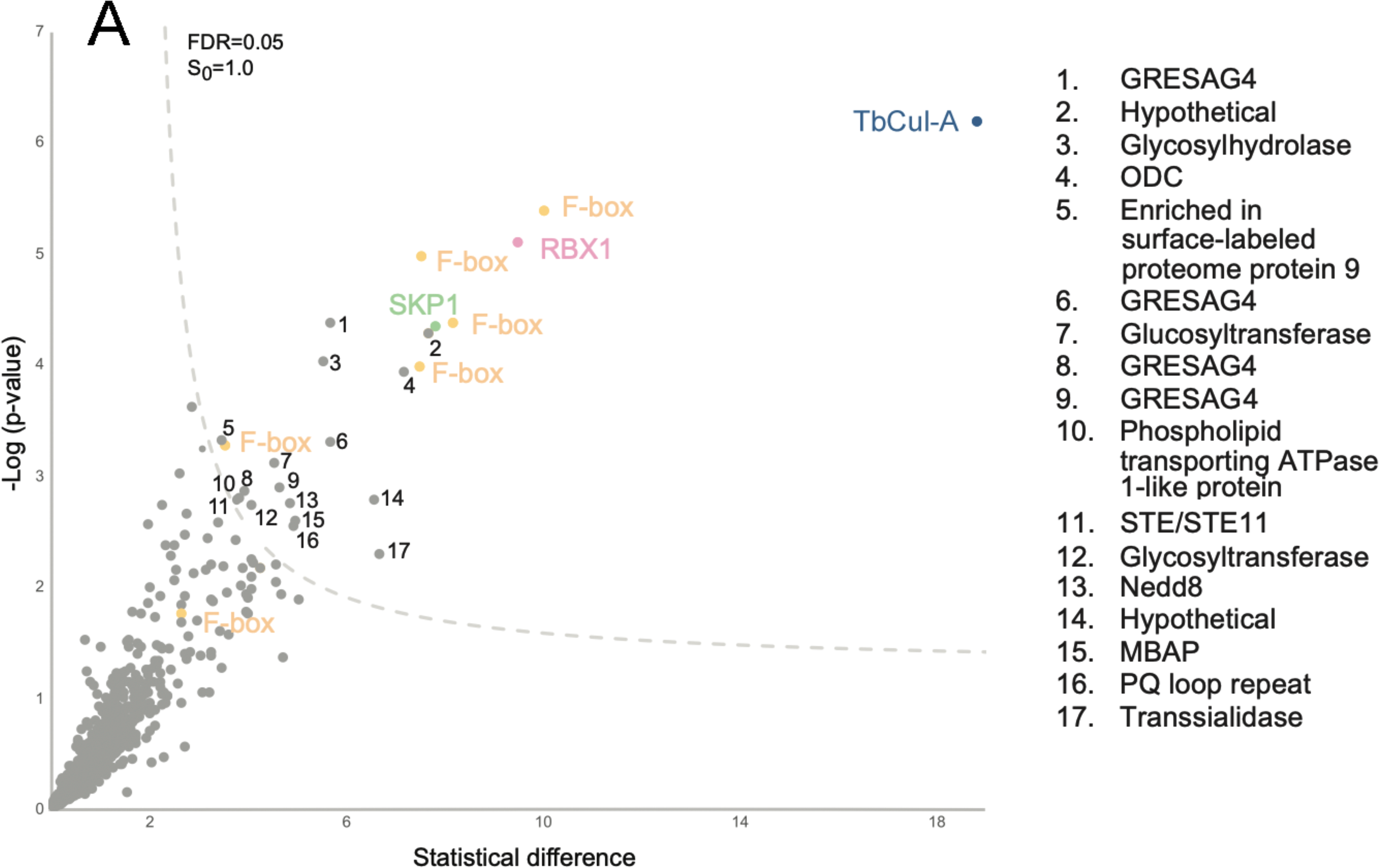
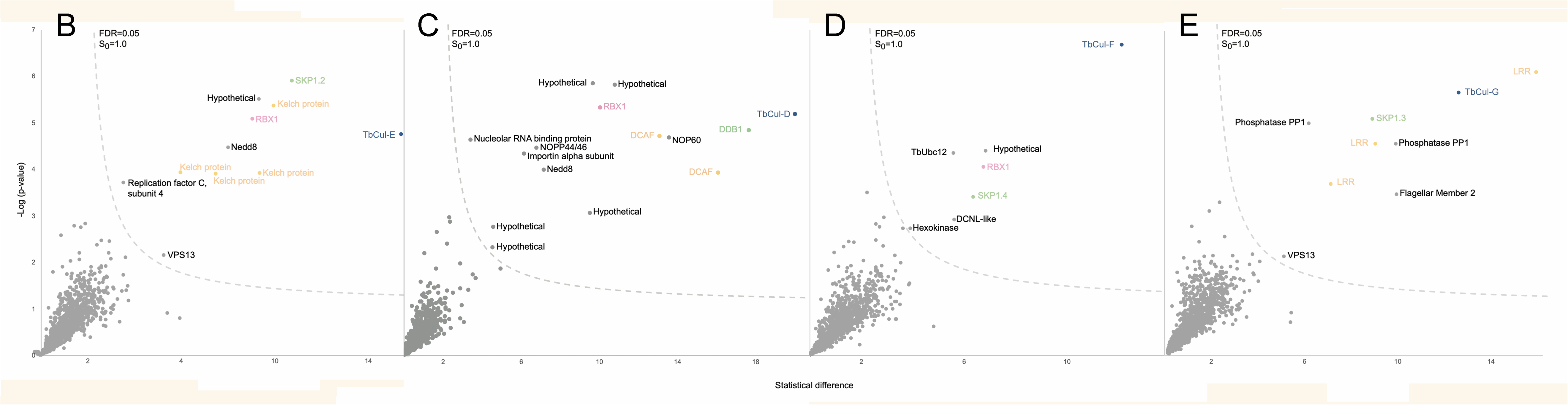
Volcano plot of proteins that co-eluted with TbCul immunoprecipitations. Panel A: Proteins that co-eluted with the HA-tagged TbCul-A after immunoprecipitation using magnetic nanobeads were identified using mass spectrometry. On the x-axis, the statistical t-test difference of LFQ intensities in the tagged cell line versus the parental is plotted and –log10p-value on the y-axis (n >3). Cutoff curves were defined for a false discovery rate of 5% and minimum fold change S_0_ at 1.0. Each protein identified is colour coded based on the expected role in the Cullin-RING complex. Panels B to E: As for panel A but for proteins coenriched with TbCul-E, D, F and G respectively. GRESAG, gene related to expression site associated gene; ODC, ornithine decarboxylase; MBAP, membrane bound acid phosphatase, DDB1, DNA damage-binding protein 1; DCAF, DDB1 and cullin associated factor; NOP60, nucleolar protein 60.

**Table 1:** Summary of protein interactions identified from cullin immunoisolations. Accession numbers for interring proteins identified that could be classified into RING, SKP or adaptor classes are given. Full details of LC-MS/MS data are available in the supplementary data. Adaptors are classified based on the presence of a specific identifiable domain or architecture. 1; No transfectants obtained. 2; No adaptors identified. 3: No RING-domain ligase identified.

| Cullin paralog | RING paralog | SKP paralog | Substrate adaptors |
| --- | --- | --- | --- |
| TbCul-A Tb927.8.5970 | RBX1 Tb927.8.5910 | SKP1.1 Tb927.11.6130 | F-box Tb927.8.1380;Tb927.5.700;cTb927.7.4300;Tb927.10.12060;cTb927.10.410; Tb927.4.3000 |
| TbCul-B<sup>1</sup> Tb927.10.7490 |  |  |  |
| TbCul-C<sup>2</sup> Tb927.8.5210 | RBX-like Tb927.5.3740 |  |  |
| TbCul-D Tb927.3.1290 | RBX1 Tb927.8.5910 | DDB1 Tb927.6.5110 | DCAF Tb927.11.3190; Tb927.8.840 |
| TbCul-E Tb927.11.11430 | RBX1 Tb927.8.5910 | SKP1.2 Tb927.9.8370 | Kelch Tb927.8.6140; Tb927.11.11450;Tb927.5.1980; Tb927.1.820 |
| TbCul-F Tb927.10.6930 | RBX1 Tb927.8.5910 | SKP1.4 Tb927.10.14310 |  |
| TbCul-G<sup>3</sup> Tb927.4.4760 |  | SKP1.3 Tb927.11.13330 | LRR Tb927.4.3100; Tb927.6.1490; Tb927.10.10190 |

### TbCul-A exhibits conserved composition

Five F-box proteins were significantly enriched in TbCul-A immunoisolations, alongside orthologs of Rbx1 and Skp1, which in *H. sapiens* comprise the archetypal SCF complex. We also identified Nedd8, two glycosyltransferases, a glycosylhydrolase, four GRESAG4 proteins, ornithine decarboxylase (ODC), a phospholipid-transporting ATPase, a kinase, a phosphatase, a *trans*-sialidase, ESAG9 and two hypothetical proteins (**Figure 4**). While the extent to which these additional proteins form a complex with trypanosome SCF is unclear, Nedd8 has been identified in *T. brucei* cullins (Liao et al., 2016), and suggests that Nedd8 controls activation of trypanosome cullin activity through conformational change (Baek et al., 2020). ODC, a key metabolic enzyme in the essential polyamide biosynthetic pathway, is the target of the human African trypanosomiasis frontline drug eflornithine that covalently binds to the ODC active site. Of the remaining, *trans*-sialidases, GRESAGs and ESAGs are all products of large paralog gene families and frequently identified in proteomics in *T. brucei,* and are hence probably not significant.

### Trypanosome cullins interact with lineage-specific subunits

We failed to identify putative interactions for TbCul-C. Among the most abundant proteins associated with TbCul-D were two proteins with DCAF domains and corresponding adaptor DDB1 and Rbx1 (Figure 4). Further, Nedd8 was detected, together with five hypothetical proteins, nucleolar proteins 44/46 and 60, a nucleolar RNA-binding protein and importin-α (O’Reilly et al., 2011). The majority of TbCul-D interactors have evidence for a nuclear localisation, including TbCul-D itself, as based on high throughput tagging, the presence of an NLS and annotation. For example,Tb927.8.1230 and Tb927.6.650 possess a nucleoplasmin-like fold and localise to the nucleolus or a DNA damage-binding protein component respectively. *H. sapiens* CUL4 includes RBX1, DDB1 and DCAF domain proteins and has roles in DNA damage repair, is located to the cell nucleus and interacts with other nuclear proteins. Hence, HsCUL4 and TbCul-D occupy a similar functional space, as well as having similar compositions.

TbCul-E, TbCul-F and TbCul-G immunoisolations did not contain F-box, DCAF or BTB proteins. This, in addition to novel Skp1 paralogs and protein families with domains characteristic of client receptors, supports TbCul-E, TbCul-F and TbCul-G as lineage-specific (Table 1). The most abundant group of proteins associated with TbCul-E were four proteins possessing three to six kelch repeats. Kelch domains span a ∼50 amino acid β-propeller structure frequently involved in protein-protein interactions and are often present in proteins also bearing F-box and BTB domains, but F-box or BTB domains are absent from TbCul-E interactions. Significantly, several kelch domain-containing proteins including KLHDC2, KLHDC3, and KLHDC10 interact with the Cullin2-RING (CRL2) complex (Bennett et al., 2010; Mahrour et al., 2008). Other interactors of TbCul-E include a Skp1 paralog, Skp1.2. Furthermore, Nedd8, replication factor C subunit four and vacuolar protein sorting protein 13 (Vps13) were detected (**Figure 4**). Excepting Nedd8, the physiological relevance of these putative protein protein interactions is unclear.

TbCul-F proved challenging due to low expression (Figure S1), but we were able to identify interactors including TbRbx1 and a further Skp1 paralog, designated TbSkp1.4 (Tb927.10.14310). TbCul-F is the sole *T. brucei* cullin for which there is no direct evidence for neddylation (Liao et al., 2016), but as E2 and E3 neddylation cascade enzymes were identified, specifically TbUbc12 and a DCNL-like protein respectively, it is likely that TbCul-F is neddylated. Another TbCul-F interaction partner is hexokinase; significantly in *H. sapiens* hexokinase is ubiquitinated by Perkin E3 ligase, but as trypanosomes lack a Perkin, ubiquitylation of trypanosome hexokinase cannot use this mechanism and hence regulation may be mediated *via* TbCul-F.

TbCul-G interacts with Skp1 paralog TbSkp1.3 (Tb927.10.13330) (Table 1) and three leucine-rich repeat proteins (LRR). LRR domains are frequently involved in protein-protein interactions and present in some F-box proteins in metazoa, making it highly likely that these LRR proteins are client recognition subunits. Further, three hypothetical proteins are also enriched, Tb924.4.3620, Tb927.4.3640 and Tb927.10.7290. The former two are phosphatases and a recent tandem duplication with near identical sequence. Tb924.4.3620 is clearly localised to the flagellum by high throughput analysis (Billington et al., 2023). The third protein, Tb927.10.7290 is FLAM2, a component of the flagellum/PFR assembly (Subota et al., 2013). As this cohort of proteins are concordant, we suggest that TbCul-G possesses novel LRR-domain client recognition subunits with roles in modulating flagellar function and/or assembly.

### Diversity of cullin client protein adaptors in trypanosomes

Some lineage-specific cullin complexes have evolved client adaptors beyond canonical proteins, for example animal-specific Cul2 and Cul5 belong to the Culα clade but recruit EloB and EloC as adaptors. In most eukaryotes where Cul2 and Cul5 are absent, EloB and C regulate tRNA polymerase II, presumably an example of repurposing (Okumura et al., 2012). EloB has a Skp1-like domain similar to some proteins identified in immunoisolates from TbCul-E, TbCul-F and TbCul-G. To investigate the origins of these proteins we constructed a phylogeny across eukaryotes, using EloB as outgroup (**Figure S2**). TbSkp1 belongs to a clade with members from every supergroup, including *bona fide* Skp1 orthologs in all sampled lineages. Significantly, TbSkp1.2, TbSkp1.3 and TbSkp1.4 are from clades restricted to Discoba; TbSkp1.2 is present only in Euglenozoa while TbSkp1.3 and TbSkp1.4 are restricted to trypanosomatids.

Given this expansion we questioned whether the RBX family has undergone similar events. Orthology searches spanning eighteen species representing all supergroups were performed using RBX1 and RBX2 orthologs and APC subunit 11 as outgroup (**Figure 5**). This resulted in the identification of three Rbx-like proteins in Discoba; only the RBX1 ortholog was identified in TbCul-A, –D, –E and –F immunoisolations. The two apparent cullin-independent Rbx-like proteins, Tb927.5.3740 and Tb927.5.3745 are a trypanosome-specific expansion and the result of a recent duplication (**Figure S6**); while Tb927.5.3745 is essential, both are of unknown function and were not present in any cullin immunoisolation.

**Figure 5:**
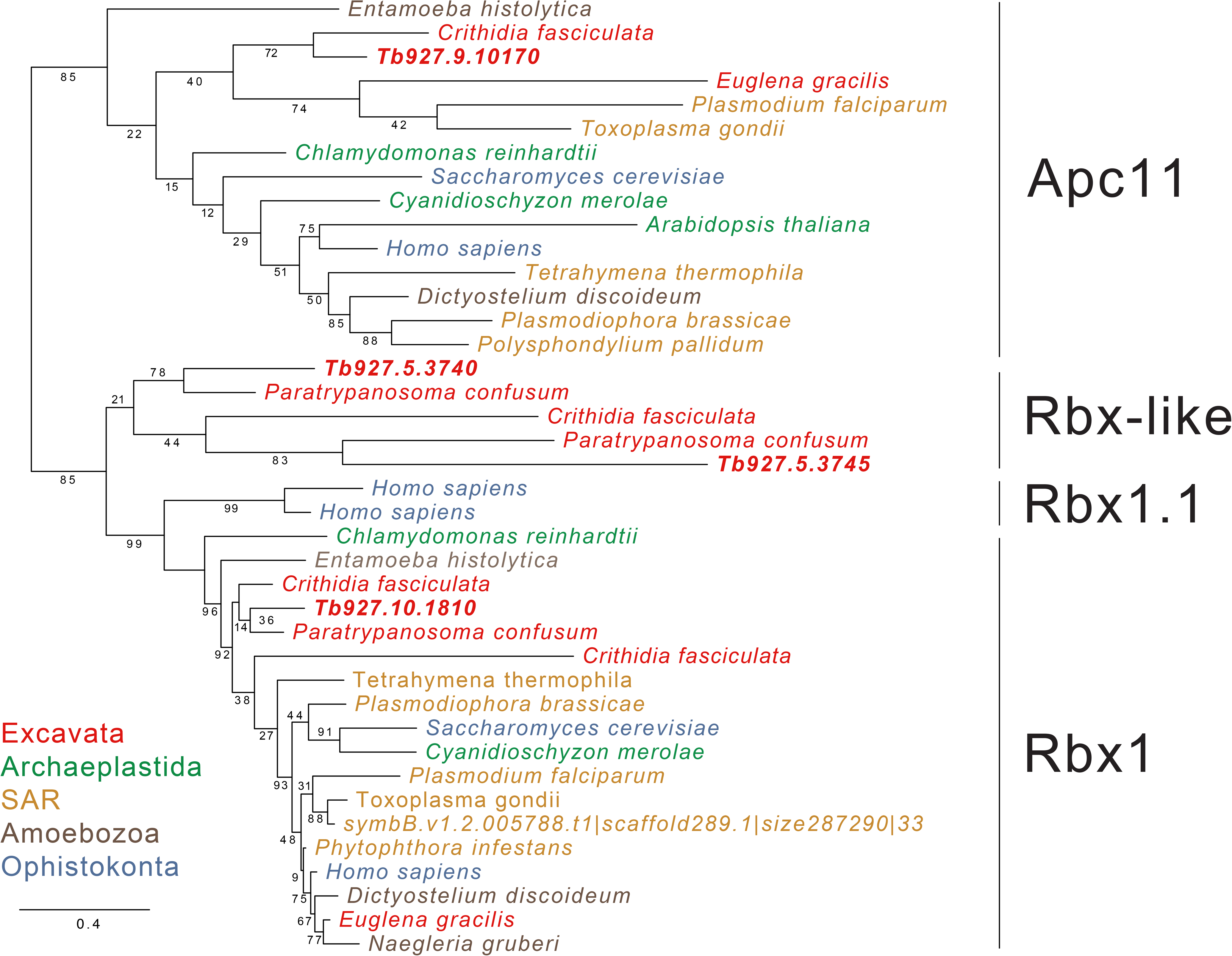
Two new RBX-like proteins in Excavata. Phylogenetic tree of RBX proteins in Eukaryota indicates a unique expansion in Kinetoplastida, that have an RBX1 ortholog and two novel RBX-like proteins. The clades identified are labelled with black bars and each taxon is coloured based on the supergroup it belongs to. The Anaphase Promoting Complex subunit 11 (APC11) was used as an outgroup to root the tree. Statistical support is indicated for PhyML/MrBayes.

### Excavata client recognition subunits

There are over 70 F-box proteins in *H. sapiens,* each presumed to recruit proteins for ubiquitylation by CUL1 (Reitsma et al., 2017). For complexes formed with CUL4, there are nearly 90 DCAF proteins (Jackson and Xiong 2009). We sought to determine if there was correspondence for F-box or DCAF proteins associated with TbCul-A and TbCul-D.

F-box protein sequences from *H. sapiens* and *A. thaliana* failed to identify a *bona fide T. brucei* F-box protein, but using “F-box” as text term at tritryp.DB followed by validation by sequence analysis using HMMER, InterPro and SMART we identified a cohort of 16, which contained all F-box proteins associated with TbCul-A. These 16 *T. brucei* F-box protein sequences were used as queries in an orthology BLASTp search against a selection of Kinetoplastida (**Figure S3**). Of the TbCul-A F-box proteins, one is restricted to African trypanosomes and close relatives such as *T. grayi* (Tb927.10.12060) and five have orthologs in *T. cruzi* (Tb927.7.4300; Tb927.8.1380; Tb927.5.700; Tb927.10.410; Tb927.4.3000) of which two are also present in *Leishmania* (Tb927.10.410; Tb927.4.3000). In addition to expression of all six in the procyclic stage of *T. brucei*, proteomics confirms expression of F-box proteins Tb927.10.410 and Tb927.4.3000 in bloodstream form *T. brucei* (tritryp.db).

Searches using 27 human *bona fide* DCAF proteins retrieved three hits from *T. brucei* (Tb927.8.4210, Tb927.9.11250 and Tb927.9.9090) as orthologs of HsWDR15b, HsDCAF13 and HsDCAF7 respectively. These, in addition to DCAFs identified following immunoprecipitation of TbCul-4 (Tb927.11.3190 and Tb927.8.840) were used as queries to reconstruct a phylogenetic tree for this family of substrate adaptors in Kinetoplastida (**Figure S5**). Every DCAF identified in *T. brucei* has orthologs in *T. cruzi* and *Leishmania* spp. except for Tb927.8.840, which is restricted to *Trypanosoma*. In addition to evidence of interaction with the cullin-RING system and a presence across Trypanosomatida, proteomics also indicates expression of these DCAF proteins in procyclic and bloodstream stages of *T. brucei* (tritryp.db).

### Knockdown of selected cullin complexes confirms specificity and interactions

To investigate trypanosome cullin functions we selected TbCul-A and TbCul-E; TbCul-A interacts with ODC, the target of the suicide-inhibitor eflornithine, and which remains in use for treatment of trypanosomiasis (Kasozi et al., 2022), while TbCul-E was considered based on essentiality and the possession of novel substrate adaptors (Alsford et al., 2009). Previous studies focusing on cullin complex components have targeted Skp1 or specific client adaptor proteins (Rojas et al., 2017, Hu et al., 2017), which in the former is likely to impact the functioning of multiple complexes and neither study investigated the impact on the cellular proteome.

Cell lines were created for inducible knockdown using the pRPaISL plasmid and quantitative reverse transcription PCR (qRT-PCR) to assess cullin transcript level after induction of RNAi for 24 or 36 hours (TbCul-E and TbCul-A, respectively); these were decreased by over 75% in both cases (**Figure S7**). Stable isotope labelling by amino acids in cell culture (SILAC) coupled to LC-MS/MS was used to quantify the effects of cullin silencing on the global protein landscape.

TbCul-A silencing had no detectable impact on proliferation, as previously reported (Rojas et al., 2017). However, we found decreased choline carnitine o-acetyltransferase (CRAT), ER oxidoreductin 1 (ERO1), quiescin sulfhydryl oxidase (QSOX) and two lineage-specific hypothetical proteins which are likely axoneme-associated (Tb927.4.2920 and Tb927.11.7520) (**Figure 6)**. In animals, fungi and vascular plants, QSOX and ERO1 are components of the ER oxidative protein folding system, facilitating disulphide bond formation (Urade 2019). CRAT is involved in fatty acid biosynthesis and also regulates levels of acyl-CoA and CoA levels (Kowacs et al., 2012). Considering that none of these proteins were identified in the immunoprecipitation of TbCul-A their decreased abundance following RNAi could be secondary.

**Figure 6:**
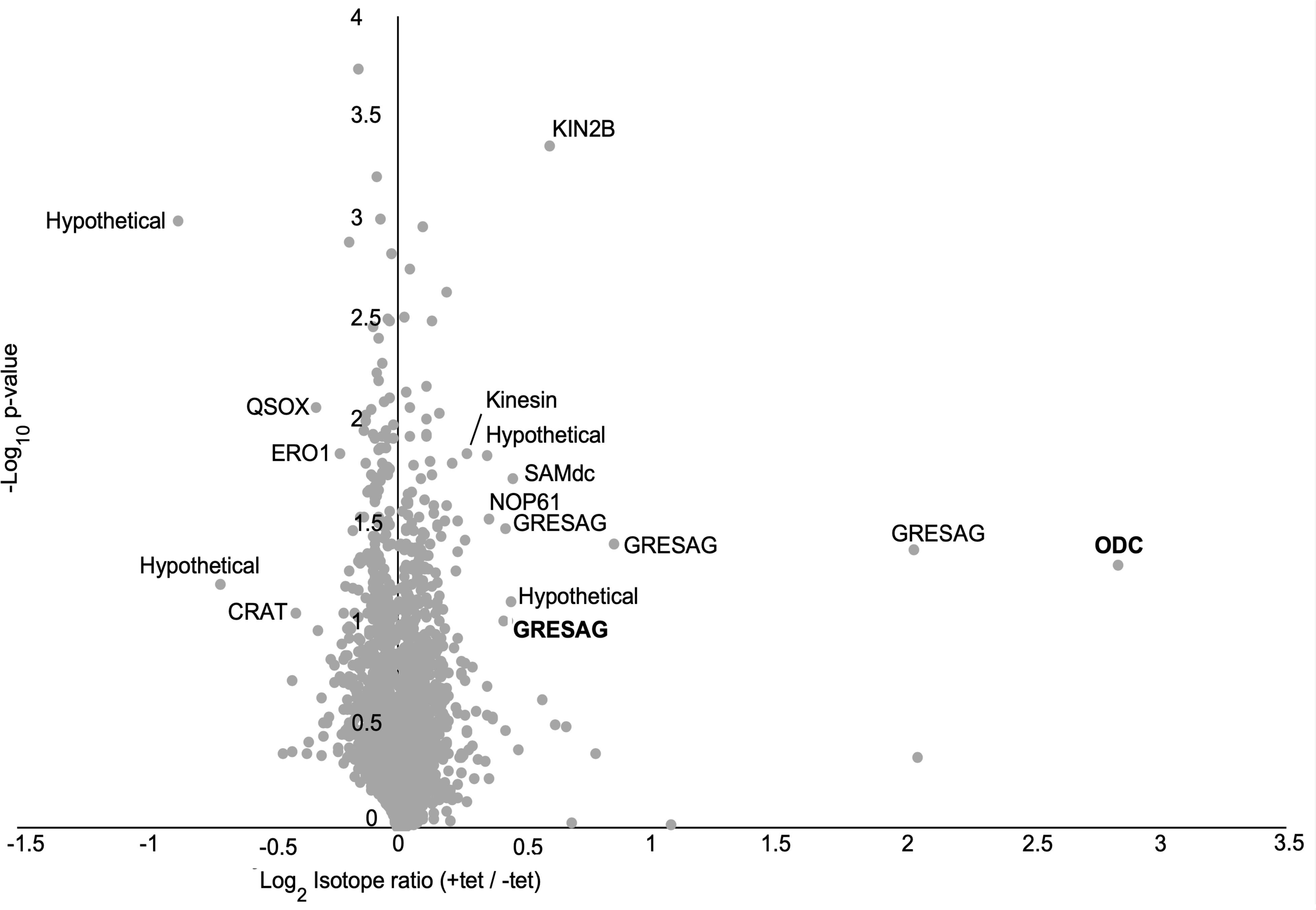
Knockdown of TbCul-A increases TbODC abundance. Landscape of *T. brucei* proteome twenty-four hours after RNAi of TbCul-A. Log_2_ transformed SILAC ratios between induced and uninduced RNAi samples (each n=3) plotted against transformed –log_10_ p-value of two sample t-test. Most affected proteins are labelled; Proteins also identified in the immunoisolates of TbCul-A are in bold. QSOX, quiescin sulfhydryl oxidase; CRAT, choline carnitine o-acetyltransferase; ERO1, endoplasmic reticulum oxidoreductin 1; KIN2B, Kinase II B; SAMdc, S-adenosyl decarboxylase; NOP61, nucleolar protein 61; GRESAG, gene related to expression site associated genes; ODC, ornithine decarboxylase.

By contrast, there was concordance between proteins with increased abundance and TbCul-A interactors upon silencing, suggesting that these proteins are TbCul-A clients. Most prominent were ODC and GRESAG paralogs, both of which interact with TbCul-A. Also impacted were S-adenosyl decarboxylase which, together with ODC, is a component of the polyamine biosynthesis pathway, two kinesins, two hypothetical proteins that are restricted to the kinetoplastida, Tb927.11.10380, which contains a RIN-domain which is associated with F-box domains in some proteins, and Tb927.10.11660, a nuclear protein with an alphafold predicted VSG-related helical bundle architecture and nucleolar protein 61 (NOP61).

To establish that ODC is a TbCul-A client protein, the ODC gene was endogenously tagged at the C-terminus with ten copies of the Ty1 epitope in the TbCul-A RNAi cell line. To determine if TbODC is ubiquitylated we used affinity isolation from whole cell lysates of cells with and without TbCul-A silenced and probed with anti-Ty1 antibody (**Figure 7**). ODC::10×Ty was detected in eluates from ubiquitin-recognising beads both before and after knockdown of TbCul-A as a single band at ∼70kDa, consistent with ubiquitylation. Moreover the signal was more intense in TbCul-A silenced cells, while RT-qPCR indicated that TbODC mRNA levels were not increased following knockdown. Further, turnover of ODC is MG132-sensitive, indicating a proteasome-dependant mechanism, also consistent with ODC as a TbCul-A substrate.

**Figure 7:**
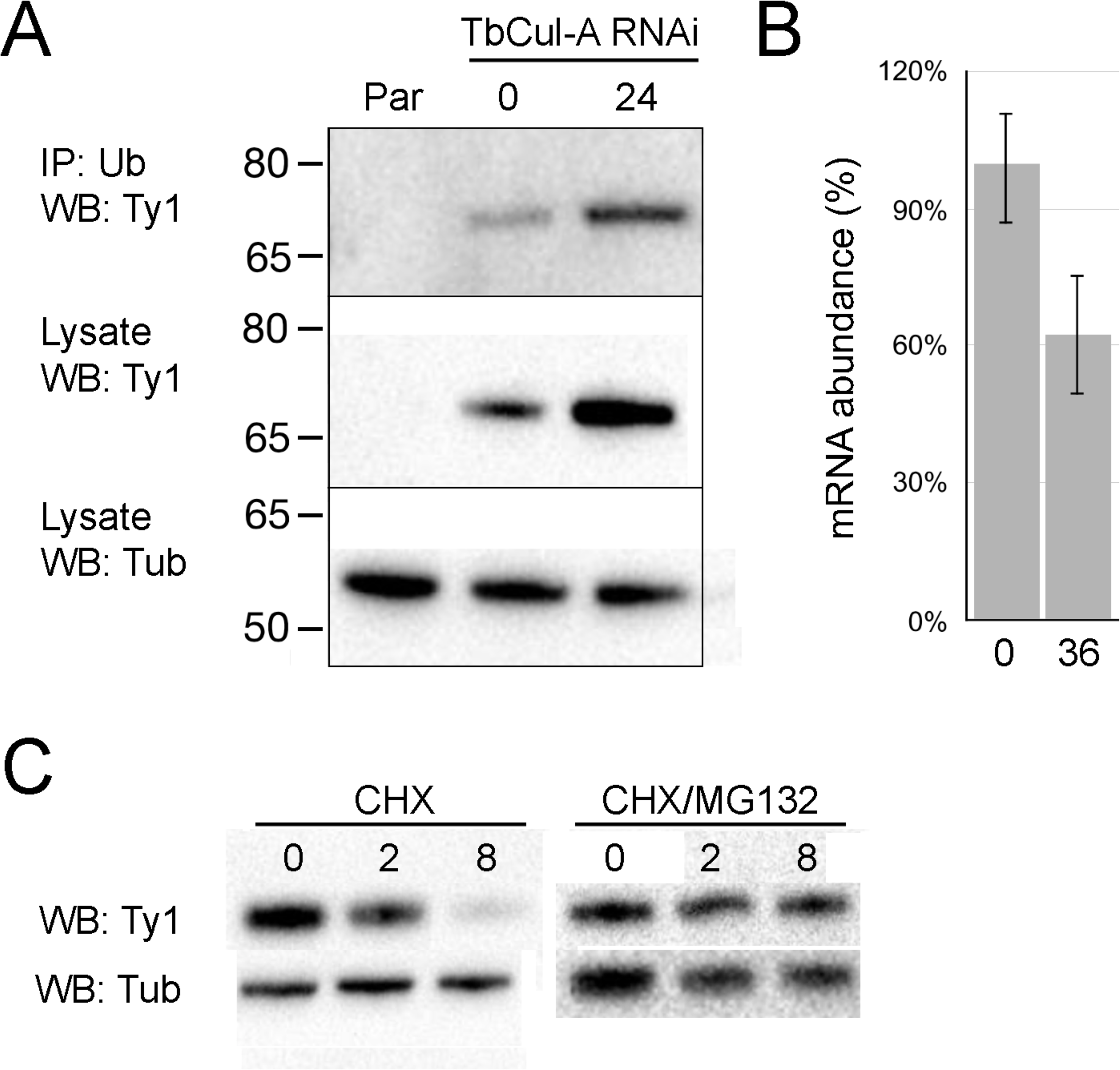
TbODC turnover is mediated by ubiquitylation and proteasome activity. Panel A: Elution of TbODC from ubiquitin-recognising beads. TbODC with a Ty1 tag endogenously fused to its C-terminus was captured by UbiQapture-Q beads after incubation with cell lysate. The intensity of TbODC-10×Ty1 detected from the beads immunoprecipitation and from the total fraction increased following induction of TbCul-A RNAi for 24 hours with tetracycline but quantitative RT-PCR (Panel B) indicates that levels of TbODC mRNA are not increased following RNAi after 36 hours. Panel C: Degradation of TbODC is mediated by the proteasome. Left; turnover of TbODC as measured following inhibition of protein synthesis with cyclohexamide (CHX) and right; turnover of TbODC as measured following inhibition of protein synthesis and proteasome activity. Blot is representative of four replicates. Par; parental T. b. b. cell line. Numbers above/below lanes indicate time post RNAi induction in hours. CHX; cyclohexamide.

Knockdown of TbCul-E is lethal (**Figure S8**), with an increased proportion of cells possessing one kinetoplast and two nuclei over time (1K2N) (**Figure 8**), broadly consistent with an earlier report (Hu et al., 2017). Replication of the kinetoplast is concurrent with development of a second flagellum, one of the earliest events indicating progression from G_1_ into cytokinesis and precedes nuclear division; the 1K2N karyotype is not part of the canonical cell cycle. Given the absence of detectable 1K0N cytoplasts, it is unlikely that these 1K2N cells arise from defective cytokinesis segregation and more likely results from disrupted signaling between replicating nuclei and kinetoplasts. A 1K2N phenotype also resulted from silencing ER-localised ERAP proteins (Wang et al 2010).

**Figure 8:**
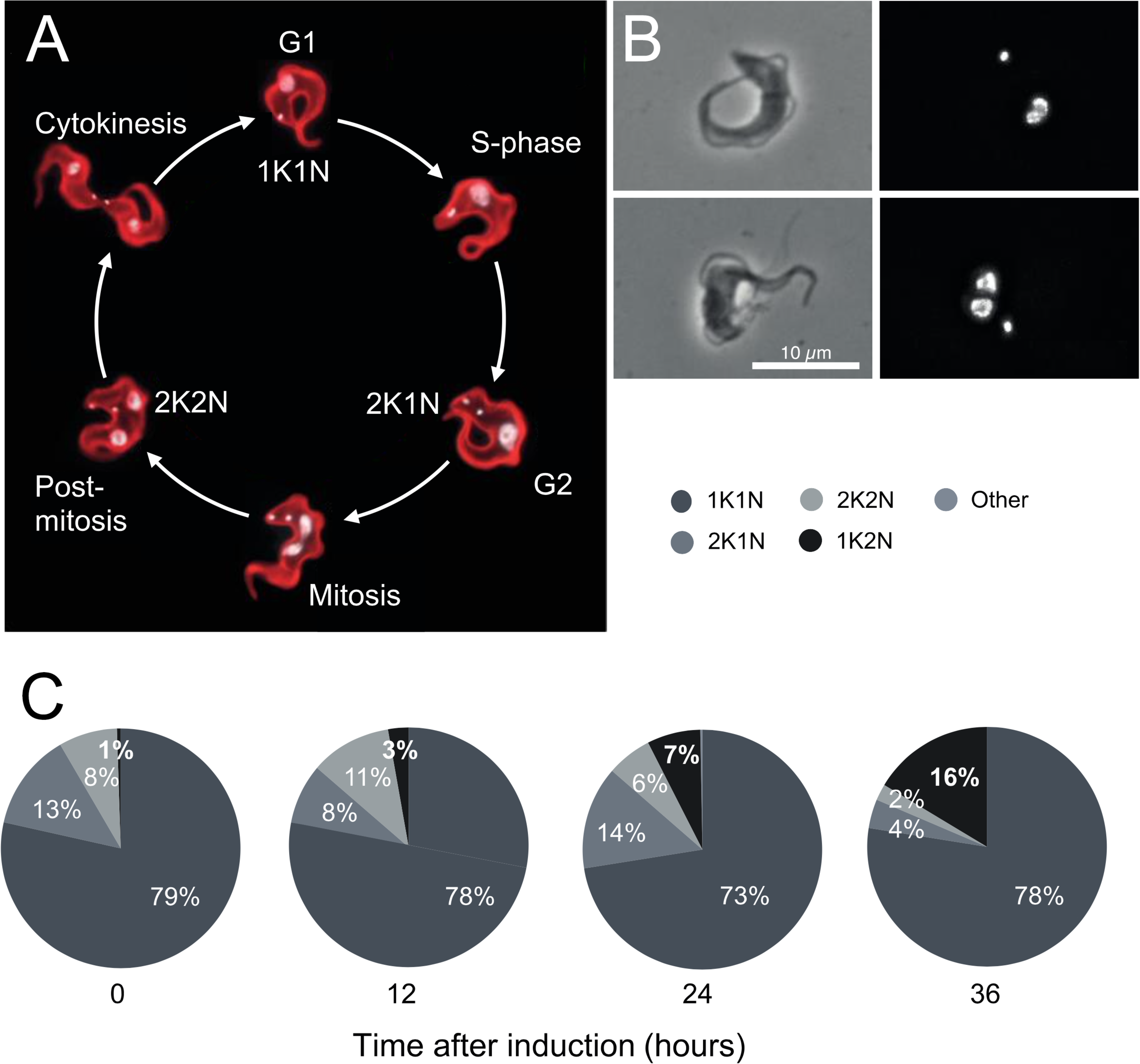
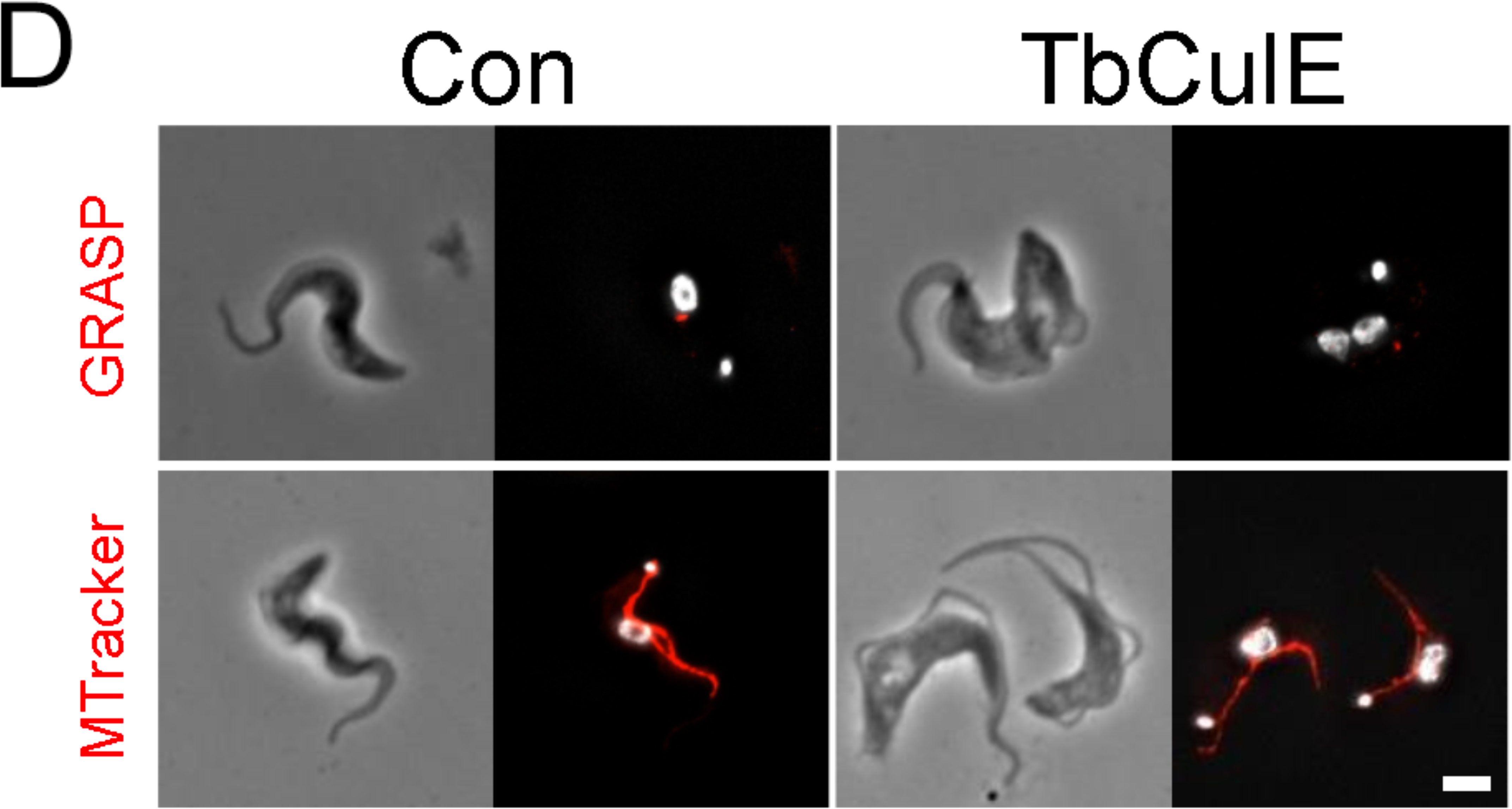
1K2N phenotype produced by knockdown of TbCul-E. Panel A; Cell cycle of bloodstream form *T. brucei* and the corresponding number of kinetoplasts and nuclei in each stage (adapted from Glover *et al*., 2019). Panel B; Example cells with one kinetoplast and two nuclei (1K2N) observed after induction of TbCUL-E RNAi. Panel C; karyotype progression throughout the knockdown (number of cells counted >300 at each time point). Panel D; Imunnofluorescence of TbGRASP and mitotracker, indicating that in 1K2N cells the Golgi complex also failed to replicate.

To interrogate the status of organelle division during cell division in TbCul-E silenced cells we used immunofluorescence against several marker proteins. A defect in timing of Golgi apparatus division was noted, as 1K2N cells also contain a single Golgi apparatus; division of the Golgi complex precedes nuclear division, and hence this suggests a general defect in coordination of division events. Analysis of the whole cell proteome following 24 hours of TbCul-E silencing (**Figure 9**) revealed multiple changes. Proteins increased in silenced cells, and hence likely TbCul-E clients are Tb927.5.2120, Tb927.10.4550, Tb927.4.5160 and kinetoplastid kinetochore protein 9 (KKT9); all are specific to kinetoplastids (Akiyoshi and Gull 2014), and consistent with the presence of novel kelch-domain client adaptors. There is some evidence for association with the basal body for Tb927.5.2120, Tb927.10.4550 and Tb927.4.5160, while KKT9 silencing is also linked to a 1K2N phenotype (Akiyoshi and Gull 2014). Further, Tb927.10.8810, a prominent down-regulated protein in TbCul-E silenced cells, is basal body localised and kinetoplastid-specific. Finally, we note that Tb927.10.4550 has a significant role in fitness (Alsford et al., 2009) and which may explain the severe loss of fitness for TbCul-E knockdown cells. Hence TbCul-E appears to mediate the stability and functions of a cohort of basal body/kinetochore-associated proteins involved in coordination of kinetoplastid-specific organelle replication events. These findings are also consistent with TbPLK, a regulator of organelle replication and where knockdown of WDR1, a client adaptor, leads to defects in basal body and bilobe replication (Hu et al., 2017).

**Figure 9:**
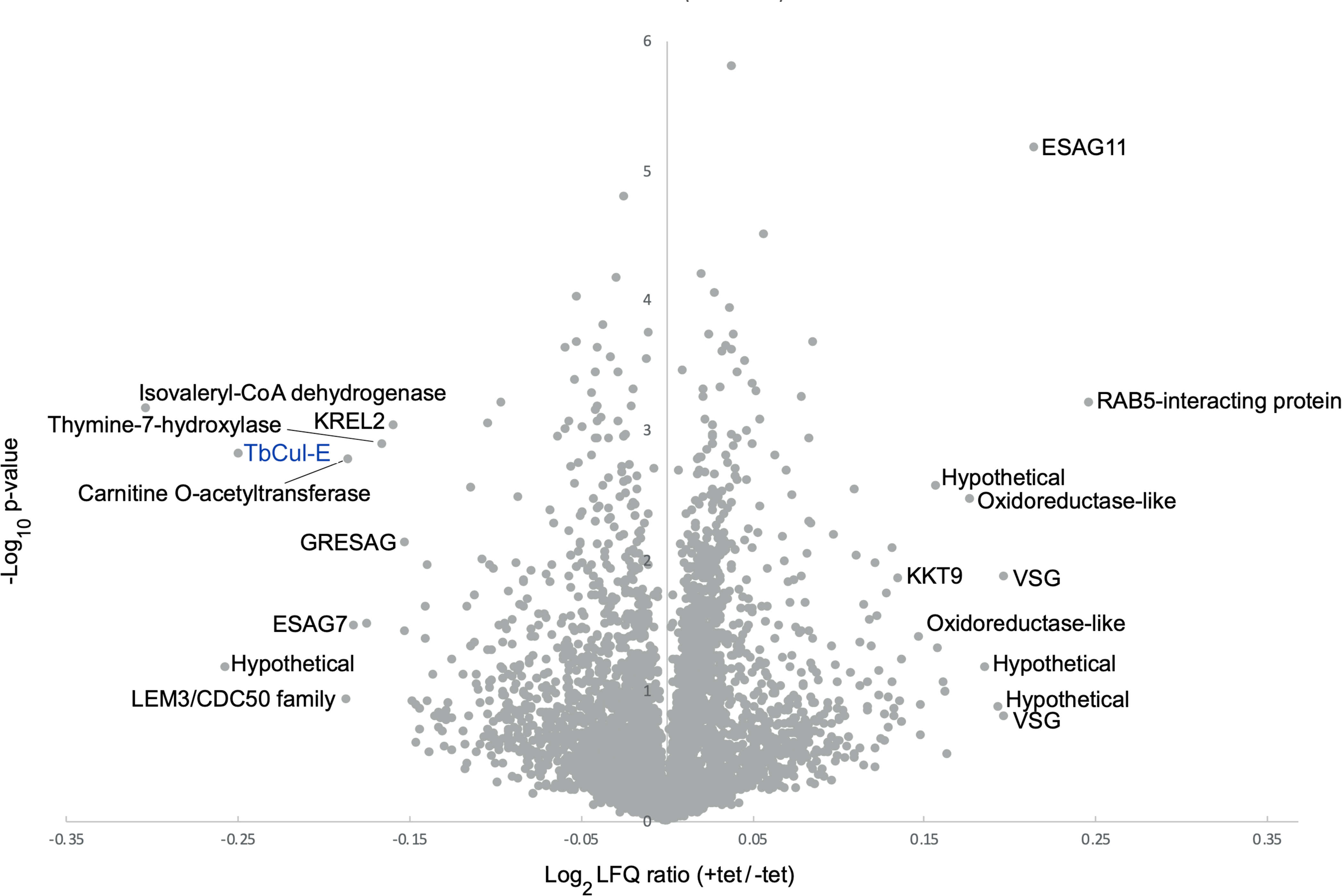
Impact of TbCul-E knockdown on global proteome. Label-free quantification of whole cell proteome after RNAi of TbCul-E after 24 hours. Log_2_ transformed label-free quantification ratios of induced/uninduced RNAi plotted on the x-axis versus –log_10_ p-value of two sample t-test on the y-axis. Data are combined from three replicates. ESAG3, expression site-associated gene 3; ZC3H47, Zinc finger CCCH domain-containing protein 47; ISG65, invariant surface glycoprotein 65kDa; PDF1, peptide deformylase 1; PGK, phosphoglycerate kinase; VSG, variant surface glycoprotein; KREL2, kinetoplast RNA-editing ligase 2; KKT9, kinetoplastid kinetochore protein 9

## Discussion

Ubiquitylation is both ancient and a central process, and in eukaryotes mediates coordination of mitosis, development, signal transduction and many other functions. As ubiquitylation connects multiple cellular systems, it is unsurprising that the machinery encompasses both conserved and lineage-specific features, reflecting a requirement to integrate both ancient/core activities with functions specific to a given lineage. At some level ubiquitylation reflects the complexity of cellular evolution; understanding the manner in which ubiquitylation varies between eukaryotic lineages is essential for uncovering such connections. To address this, *albeit* in a limited manner, we undertook combined *in silico* and experimental analysis, using the African trypanosome as a taxonomically divergent organism and selecting the cullin ubiquitin ligases for detailed analysis.

We reconstructed the evolutionary history of multiple components of the cullin complexes, including the cullin scaffold, the Rbx E3 ligase and client adaptors, revealing novel and unexpected complexity. Firstly, we showed the three previously recognised ancestral cullin clades (Culα, Culβ and Culγ) are present in SAR, suggesting a more ancient origin than previously noted. We also identified an additional clade, Culδ, which is restricted to the amorphea. Further, this new topology allows us to propose that Culα split from the remaining cullins first, followed by Culβ and Culγ, while Culδ is likely a post-LECA acquisition. While the Culα clade possesses representatives from across eukaryotes, we find that Discoba sequences are absent from Culβ, Culγ and Culδ, and instead form a unique clade we term Culκ. Despite challenging this topology through removal of sequences and recalculating, the Culκ clade was retained; while we cannot exclude a long branch attraction artefact, suggested by similar architectures of client adaptors identified for TbCul-D and human CUL4, we consider this evidence for extreme divergence regardless of descent and that Culκ provides evidence for the likely independent cullin expansion within kinetoplastida. Building on earlier analysis we also detected additional Skp1 subfamilies associated with cullin complexes (Yamada et al., 2023). Skp1 paralogs are mainly kinetoplastida specific, but importantly physically associated with cullin complexes, indicating conserved roles within the ubiquitylation machinery. Finally, we present evidence for two additional clades of Rbx1 related proteins, with evidence for both metazoan and kinetoplastida-specific Rbx1 clades. The mechanistic implications of multiple Skp1 and Rbx1 paralogs is unclear, but does suggest the potential of independent modulation of cullin activity, and also led to the evolution of a Skp1-containing debuiquitinase complex specific to kinetoplastids (Yamada et al., 2023).

Our evidence demonstrates diversity within the cullin E3 ligase system beyond animals and fungi, and hence likely engagement of cullin ligases with specific adaptive/ lineage-specific functionality. Reconstruction of F-box, DCAF and kelch domain-containing client adaptor distributions for the kinetoplastids suggests that our affinity isolations sampled all F-box subfamilies but only a subset of the DCAF and kelch families. Further work will be required to ascertain if these additional DCAF or kelch proteins are associated with a cullin complex, but increases the potential size and diversity of the trypanosomatid cullin ligase repertoire. Most trypanosome cullin complexes contain either nedd8 itself or components of the neddylation machinery, supporting that trypanosomes regulate their cullin ligases in a manner conserved with metazoa and consistent with an earlier report that six trypanosome cullins are neddylated (Liao et al., 2017). Moreover, we speculate that the presence of candidate client adaptor proteins possessing either kelch or LLR-domains alone, which in metazoa are incorporated into more complex F-box and BTB multi-domain adaptors, indicates an evolutionary pathway by which adaptor proteins have grown in architectural complexity.

Silencing of the cullin proteins from TbCul-A and TbCul-E complexes provides insights into specific functions. Significantly, we find that ODC is a client for TbCul-A, based on physical association with TbCul-A together with an increase in ODC abundance on TbCul-A silencing. ODC is the target of eflornithine, an important trypanocide. Eflornithine specificity has been suggested as due to differential turnover of human and trypanosome enzymes and that human ODC turns over rapidly, refreshing the host ODC pool, while the parasite enzyme has a more prolonged half-life and hence inhibition is more pronounced (Wang 1991). The present data contradict this model as trypanosome ODC is clearly an efficient ubiquitylation client and degraded rapidly *via* a proteasome-dependant route. It is quite possible that the earlier work, which utilised a heterologous expression system to analyse ODC turnover, provided an aberrant half-life due to incompatibility between trypanosome ODC and mammalian ubiquitylation and is more consistent with more recent analysis of trypanosomatid ODC turnover rates obtained directly in trypanosomes (Iten et al., 1997).

TbCul-E, which likely possesses lineage-specific keltch-domain client adaptors, has roles in organelle replication, most obviously evidenced by accumulation of aberrant 2K1N cells following silencing. Significantly, this impact extends to additional organelles, implying a general defect to replication coordination involving turnover of both basal body and kinetochore components. We suggest that the coordinated turnover of components of these two cytoskeletal organisers provides a mechanism underpinning mitosis and which confirms and extends evidence that TbPLK is a TbCul-E substrate (Hu et al., 2017, Ikeda and de Graffenried 2012, An et al., 2021) by identification of a larger cohort of factors likely important in control of flagellum replication and the kinetochore (Nerusheva and Akiyoshi 2016). We suggest that this final example represents a lineage-specific function of a cullin, incorporating novel client adaptors and which also control trypanosome-specific cell cycle events. This, together with our recent description of TUSK (Yamada et al., 2023), a deubiquitinase complex containing a divergent Skp paralog, that controls turnover of surface proteins, provides evidence for considerable diversification of ubiquitylation mechanisms in trypanosomes.

## Methods and materials

*Cell culture.* Procyclic cells (PCF) derived from *Trypanosoma brucei* subspecies *brucei* strain Lister 427 were grown in SDM-79 JRH 57453 media (Life Technologies UK) supplemented with 7.5 mg/l of hemin (Sigma-Aldrich) and 10U/10ug/ml of penicillin/ streptomycin (Thermo). Cells were maintained at densities between 5×10^5^ to 3×10^7^ cells/ ml in non-vented flasks (Starlab). Cell lines transfected with the pMOT-4H plasmid (Oberholzer et al 2006) were grown with 25 ug/ml of hygromycin B Gold (Invivogen).

### Genetic modifications

Cullin coding sequences were endogenously tagged using the PCR-based pMOT system (Oberholzer et al 2006). OneTaq Hot Start polymerase (NEB) was used to amplify the pMOT-4H plasmid template using long oligonucleotide primers (**Table S1**, ThermoFisher). The PCR products were purified using PCR purification columns (Qiagen) and used for transfection. Primer sequences are given in Table S1. PCF cells at log phase were resuspended in cytomix buffer (Chung, et al 2008) alongside 10ug of DNA. They were electroporated at 1.7 kV with three pulses of 100us with 200ms intervals in a BTX Gemini electroporator (ThermoFisher) using a 0.4cm cuvette (Bio-Rad) and immediately transferred to SDM-79 media. 25 ug/mL of hygromycin B Gold (Invitrogen) was added 24 h after transfection.

### Western blotting

Cells were harvested and resuspended in NuPAGE LDS sample buffer containing NuPAGE sample reducing agent (ThermoFisher). The lysate was homogenised using a sonicator (10 pulses, five seconds each) and run on NuPAGE 4-12% Bis-Tris protein gels (ThermoFisher). The iBlot 2 Dry Blotting System (ThermoFisher) was used to transfer proteins to a PVDF membrane. Membranes were blocked with 5% freeze-dried skim milk in Tris buffered saline (TBS) buffer supplemented with 0.1% tween 20 for half an hour before overnight incubation at 4°C with the primary antibody. After three washes with TBS of five minutes each, membranes were incubated with the secondary, HRP-conjugated antibody for one hour at room temperature prior to developing using ECL Western Blot Substrate (Pierce). Luminesence was visualised using a ChemiDoc MP Imaging System (Bio-Rad). Antibodies were used at the following dilutions: Rat anti-HA IgG1 (clone 3F10; Sigma) at 1:2000, mouse anti-Ty1 (SAB4800032, Sigma) at 1:10000, mouse β-tubulin (clone KMX-1; Millipore) at 1:10000. Secondary antibodies coupled to horseradish peroxidase (HRP) were anti-rat-HRP at 1:10000 and anti-mouse-HRP at 1:10000 (Sigma).

### Cryomilling and immunoprecipitation

Four litres of PCF *T. brucei* were grown as detailed above in 2 L roller bottles and harvested when density reached 2×10^7^ cells/ml. The cell pellet was washed with PBS supplemented with protease inhibitors (mini cOmplete cocktail, Roche) and snap frozen in liquid nitrogen. The frozen pellet was ground into a fine powder using a P100 mill (Retsch) as described (Obado et al 2016).

Fifty milligrams of cell powder were resuspended in buffer (20mM HEPES pH 7.4, 250mM NaCl, 1mM Mg_2_Cl and 0.01 mM CaCl_2_ 0.1% (w/v) Brij58). After sonication, the homogenised cell suspension was centrifuged at 4°C and 11000rpm for 10 minutes. The supernatant was incubated with Pierce anti-HA Magnetic Beads (ThermoScientific) for 2 hours at 4°C. Supernatant was discarded and the magnetic beads washed three times using resuspension buffer with a reduced detergent concentration (0.01% (w/v) Brij58). The proteins bound to the beads were eluted at 70°C for 10 minutes in NuPAGE loading buffer and supplemented with sample reducing agent (ThermoFisher).

### Silver staining

Visualisation of proteins in NuPAGE gels was performed using the SilverQuest Silver Staining kit (ThermoFisher) following the manufacturer’s protocol. Gels were submerged in fixative (40% ethanol, 10% acetic acid (v/v)) for 20 minutes before incubation for 10 minutes in sensitizing solution. Gels were washed in 30% ethanol (v/v), followed by incubation in the staining solution for 15 minutes. Before incubation in the developer solution, the gels were briefly washed with MilliQ Ultrapure water. Gels were imaged using a ChemiDoc MP Imaging System (Bio-Rad).

### RNAi with pRPaiSL

To generate tetracycline inducible RNAi cell lines, we used the pRPaiSL (Alsford and Horn, 2008) system. Briefly, the best region of the cullin genes to be targeted by the RNA stem-loop was identified using RNAit2 (Redmond, Vadivelu and Field, 2003), which also provided the appropriate oligo primers (Table S1), synthesised by Thermo Fisher. PCR products were purified and digested using the appropriate enzymes (NEB) as described by Alsford S. *et al*. (Alsford and Horn, 2008), before using T4 DNA ligase (NEB) to insert it into the linearized pRPaiSL MCS1/2 plasmid. The complete pRPaiSL with both PCR inserts was digested with AscI and the fragment of interest was cleaned-up using a gel extraction kit (Qiagen) to be used for transfection.

### RNA extraction and quantitative RT-PCR

RNA extraction was performed using the RNeasy Kit (Qiagen). Briefly, 5×10^7^ cells were harvested (800g, 24°C, 10min) and washed twice with PBS before lysis using the RLT buffer. The lysate was mixed with 70% ethanol in DEPC-treated water (Invitrogen) and processed essentially following the manufacturer indications and supplied columns. Reverse transcription of RNA and quantitative PCR was performed using the Luna Universal One-step RT-qPCR Kit (NEB) using 1 µg of RNA per reaction. Reactions were carried out in a QuantStudio 3 real time PCR system (ThermoFisher) and included reverse transcription at 55°C for 10 minutes, followed by initial denaturation at 95°C for 1 minute followed by 40 cycles of 95°C for 10 seconds and 60°C for 30 seconds with a signal read at the end of each cycle plus a final melting curve to check fidelity from 60-95°C, with a signal read every 1°C.The data was normalised against the *T. brucei β*-tubulin gene. Fold changes in gene transcription were calculated using the Ct method, normalised to the parental strain *T. brucei* 2T1 and displayed as relative quantity.

### Proteomics

Affinity capture eluates from the magnetic beads were loaded onto NuPAGE Bis-Tris 4-12% gradient polyacrylamide gels as described previously (Zoltner et al 2020). Briefly, samples were run ∼1 cm into the gel and cut using a sterile scalpel. Slices were subjected to tryptic digest and reductive alkylation using routine procedures and eluted peptides analysed by LC-MS/MS on a UltiMate 3000 RSLCnano System (Thermo Scientific) coupled to a LTQ OrbiTrap Velos Pro mass spectrometer (Thermo Scientific).

TbCul-E RNAi samples were prepared label-free at two times (12 and 24h after induction with 1ug/ml tetracyclin). Uninduced cells were used as controls. Cells were washed with PBS containing complete protease inhibitors (Roche), extracted with 1× NuPAGE sample buffer and sonicated. Lysates containing 1×107 cells were fractionated on a NuPAGE Bis-Tris 4-12% gradient polyacrylamide gel (Thermo) under reducing conditions. The approximately 30 mm migration portion was contained in three slices (fractions) that were subjected to tryptic digest and reductive alkylation using routine procedures. The eluted peptides were then analyzed by liquid chromatography-tandem mass spectrometry (LC-MS2) on a ultimate3000 nano rapid separation LC system (Dionex) coupled to a Q Exactive HF (Thermo-Fisher Scientific).

TbCul-A RNAi samples were prepared using SILAC. HMI-9 for SILAC was prepared as described previously (Zoltner et al., 2015). Either normal L-Arginine and L-Lysine (HMI11-R0K0), or L-Arginine−^13^C_6_ and L-Lysine ^4,4,5,5-2^H_4_ (HMI11-R_6_K_4_) (Cambridge Isotope Laboratories) were added at 120 µM and 240 µM respectively (Cox et al., 2008, 2014). Cells with induced TbCul-A RNAi were grown in parallel with uninduced cells, in the presence of HMI11-R_0_K_0_ or HMI11-R_6_K_4_, respectively. Cultures in logarithmic growth were mixed after 24 hours and 48 hours, immediately harvested by centrifugation, washed twice with PBS containing protease inhibitors (Roche) and resuspended in Laemmli buffer containing 1 mM dithiothreitol. Samples were generated in triplicate and one label swap was performed. Samples were sonicated and aliquots containing 5×10^6^ cells separated on a NuPAGE bis-tris 4–12% gradient polyacrylamide gel (Thermo). The sample lane was divided into eight slices that were excised from the Coomassie stained gel, destained, then subjected to tryptic digest and reductive alkylation. The eight fractions obtained from SDS-PAGE were subjected to LC-MS/MS on a UltiMate 3000 RSLCnano System (Thermo Fisher Scientific) coupled to an Q Exactive HF (Thermo Fisher Scientific) mass spectrometer.

Spectra were processed in MaxQuant V1.5. or 1.6.1.0 (Cul-E RNAi) searching the *T. brucei brucei* 927 annotated protein database (release 42) from TriTrypDB (Aslett et al., 2010). Affinity capture and Cul-E RNAi spectra were processed using the intensity-based label-free quantification (LFQ). For TbCul-A RNAi, SILAC ratios were calculated using only peptides that could be uniquely mapped to a given protein. False discovery rates (FDR) of 0.01 were calculated at the levels of peptides, proteins and modification sites based on the number of hits against the reversed sequence database. Perseus (Tyanova et al 2016) was used for statistical analysis. LFQ values were log2 transformed, and missing values imputed from a normal distribution of intensities around the detection limit of the mass spectrometer. For the affinity capture, these values were subjected to a Student’s t-test comparing an untagged control (wt) sample group to the 3×HA-tagged protein triplicate sample groups. −log10 t-test p-values were plotted versus t-test difference to generate volcano plots. Potential interactors were classified according to their position in the volcano plot, applying cutoff curves for “significant class A” (SigA; FDR = 0.01, s0 = 0.1), “significant class B” (SigB; FDR = 0.05, s0 = 1). The cutoff is based on the FDR and the artificial factor s0, controlling the relative importance of the t-test p-value and difference between means (at s0 = 0 only the p-value matters, whereas at nonzero s0, the difference of means contributes).

### Bioinformatics

*Homo sapiens* cullin proteins (Uniprot ID: Q13616, Q13617, Q13618, Q13619, Q13620, Q93034, Q14999 and Q8IWT3) were used as query against the genome of *Trypanosoma brucei brucei* TREU927 in a BLASTp (Altschul et al., 1990) search (BLOSUM62, Existence: 11 Extension: 1 Conditional compositional score matrix adjustment) to retrieve cullin genes in *T*. *brucei*. The *Trypanosoma* sp. hits were then used as query in a BLASTp against their own genome and the top 10 hits considered for further analyses. Using ScrollSaw (Elias et al 2012), the selected hits were used as queries in reverse BLASTp searches against curated lists of organisms of the Archaeplastida, SAR, Excavata, Amoebozoa and Ophistokonta. As cut-off, a maximum of five hits per organisms per query with an e-value <0.001 and coverage >30% were recorded. These hits, as well as those identified using the subunit 2 of the anaphase promoting complex (Uniprot ID: Q9UJX6) as query, were aligned and edited using MUSCLE (Edgar 2004), only allowing 25% or less of the entry sequences to have a gap in every given amino acid position. FastTree (Price et al 2009) was used to generate initial phylogenetic reconstruction of the cullin family for each of the supergroups of Eukaryota. Two protein entries per clade per tree were selected, pooled and aligned as described above to create a pan-eukaryotic cullin tree using FastTree, PhyML (Guindon et al 2005) and MrBayes (Huelsenbeck and Ronquist 2001). For PhyML, 1000 bootstrap and the LG model were used while FastTree was used with default settings. MrBayes used BEAGLE and the following block was added to the input files: nrun=2 nchains=8 lset rates=gamma Ngammacat=4 nucmodel=protein code=universal; prset aamodelpr=mixed; mcmcp ngen=8000000 relburnin=yes burninfrac=0.25 printfreq=100000 samplefreq=1000 diagnfreq=1000 nchains=8 savebrlens=yes;

Phylogenetic analysis of Skp and Rbx did not require ScrollSaw, and reverse BLASTp searches against a curated selection of eukaryotes from Archaeplastida, SAR, Excavata, Amoebozoa and Ophistokonta were used to identify orthologs and paralogs in each of these supergroups. The queries used were *Homo sapiens* SKP1, RBX1 and RBX2 as well as ELOB and APC11 as outgroups (Uniprot ID: P63208, P62877, Q9UBF6, Q15370, Q9NYG5). Other Skp and Rbx candidates identified from the immunoprecipitation of cullin complexes in *Trypanosoma brucei* were also used as query (Tb927.10.11610, Tb927.9.8570, Tb927.11.13330, Tb927.10.14310, Tb927.11.6130). Again, the cut-off was a maximum of five hits per organisms per query with an e-value < 0.001 and coverage >30% were recorded. For each of the trees, the hits were aligned and edited as described above and the phylogenetic trees were reconstructed using FastTree, PhyML and MrBayes using the same parameters as stated before. The reverse BLASTp searches across eukaryotes, the alignment of sequences and generation of trees with FastTree were performed using an in house automated Python script (https://github.com/laelbarlow/ ERB_pipeline_output_interpretation_workflow). Trees were visualised, annotated and coloured using FigTree (Rambaut, A. 2018). Fasta files for the sequences used to generate each phylogenetic tree and a list of species for each are provided as a supplementary data archive.

Reconstruction of substrate receptor families was generated using orthology BLASTp (Altschul et al., 1990) searches against a curated list of taxa with *T. brucei* proteins as query. Default settings were applied (BLOSUM62, Existence: 11 Extension: 1 Conditional compositional score matrix adjustment) and the top five hits with an e-value <0.0001 and a coverage >70% were recorded. Hits were aligned and edited using MUSCLE (Edgar 2004), only allowing 25% or less of the entry sequences to have a gap in every given amino acid position. FastTree (Price et al 2009), PhyML (Guindon et al 2005) and MrBayes (Huelsenbeck and Ronquist 2001) were used to generate the phylogenetic reconstruction. Reverse BLASTp searches using either *H. sapiens* or *A. thaliana* proteins as query were performed in the same manner.

### Immunofluorescence microscopy

*Trypanosoma brucei* cells were harvested and washed twice using Dulbecco’s PBS (Thermo), then fixed using 3% paraformaldehyde (Thermo) at 37°C for 10 minutes and resuspended in Dulbecco’s PBS. Cells were allowed to settle onto a poly-lysine slide (VWR) for approximately 45 minutes at room temperature. The cells were then permeabilized using 0.2% Triton X-100 for 10 minutes and incubated for 1 hour with 20% FBS. Primary antibodies were used to incubate the cells at 4°C overnight, followed by incubation with secondary antibodies for 1 hour at room temperature. Primary antibody concentrations were rabbit anti-GRASP (from M. Ferguson) at 1:1000 and secondary antibody (Thermo) anti-rabbit Alexa-568 at 1:1000. MitoTracker (Thermo) was added to cultures to a final concentration of 1nM 5 minutes before harvesting. Vectashield with 4,6-diamidino-2-phenylindole (DAPI; Vector Laboratories, Inc.) was applied to slides before covering them with a coverslip (WVR). Cells were visualised using a Zeiss Axiovert 200 microscope with an AxioCam camera and ZEN Pro software (Carl Zeiss, Germany) and images acquired as z-stack of 0.26µm. Images were generated using OMERO (https://www.openmicroscopy.org/omero/).

## Acknowledgements

This work was supported by grants from the Wellcome Trust (204697/Z/16/Z) and the Medical Research Council (MR/P009018/1) to MCF. We thank Lael Barlow (Dundee) for custom phylogenetics/search scripts. Proteomics data associated with these analyses have been deposited at the Pride Proteomics database at https://www.ebi.ac.uk/pride with accession number: PXD042022.

## Supplementary material for

### Supplementary figure legends

**Figure S1:**
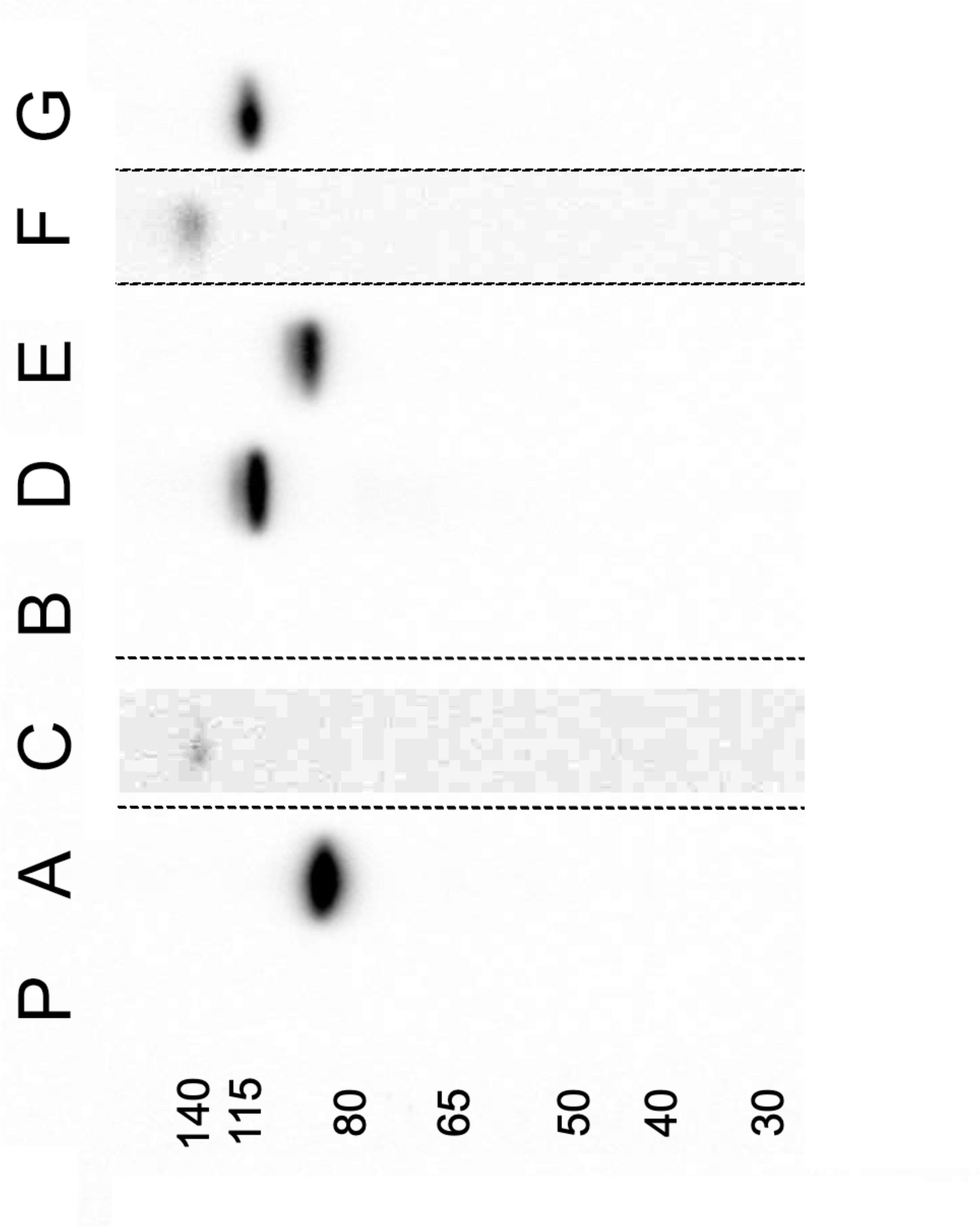
Validation and relative abundance of tagged *T. brucei* cullins. Western blotting against the human influenza hemagglutinin (HA) epitope endogenously fused to the cullin genes of *T. brucei*. Note the low abundance of TbCul-F and TbCul-C. No clones successfully expressing TbCul-B were obtained. Molecular weight markers are shown at left in kDa. All samples were separated in the same polyacrylamide gel but the chemiluminescence in the TbCul-C and TbCul-F lanes have been contrast enhanced over the regions indicated with dotted lines. P is parental line.

**Figure S2:**
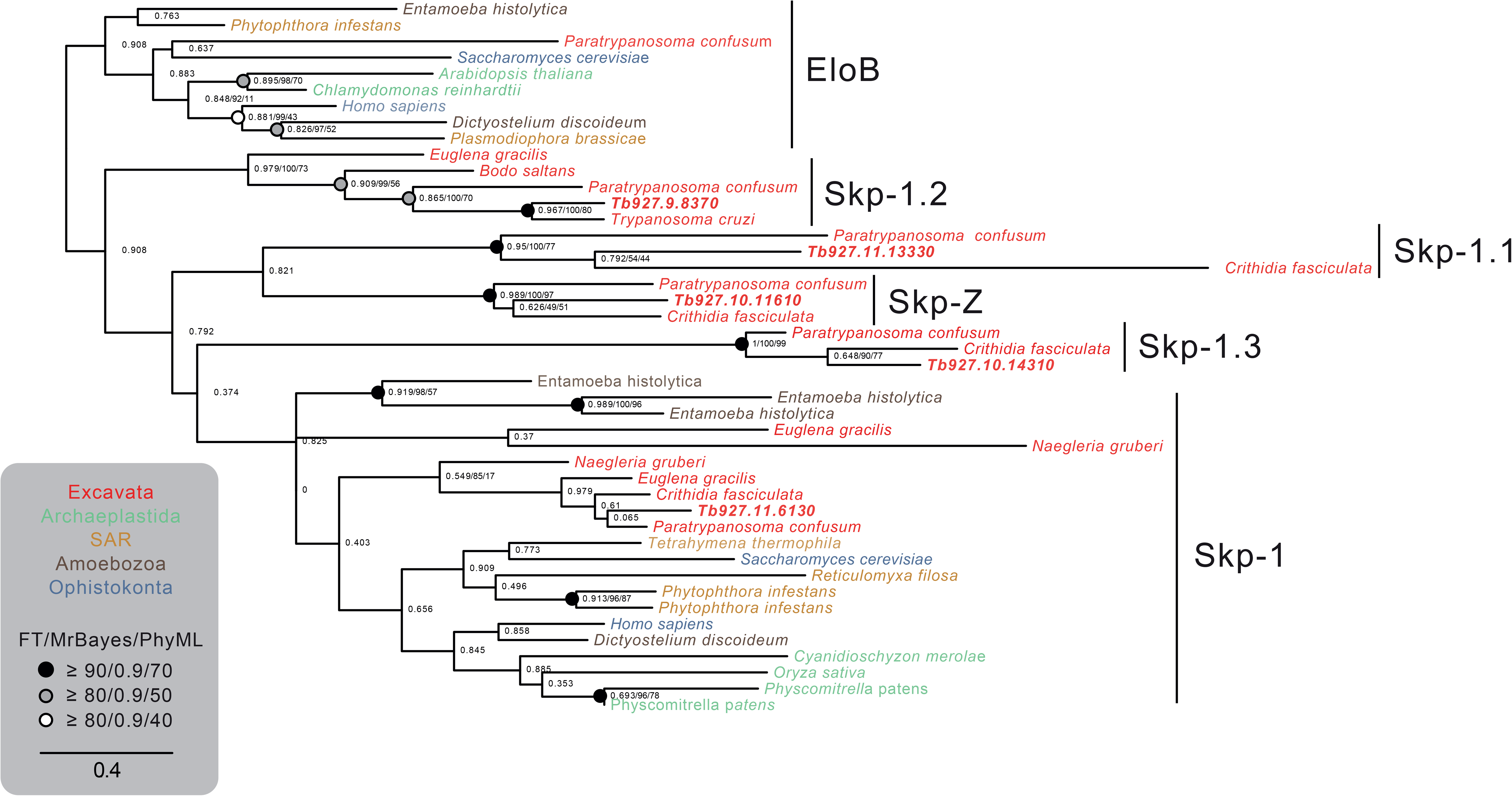
The SKP1 family is expanded in Excavata. Phylogenetic reconstruction of the SKP1 family of protein adaptors in Eukaryota. Skp1.1 is present in all eukaryotic supergroups while Excavata has four additional members: Skp1.2, Skp1.3, Skp1.4 and Skp-Z. The five clades are indicated with black bars and each taxon is coloured based on the supergroup to which it belongs. Elongin factor B (EloB), that acts as protein adaptor in some *H. sapiens* cullin-RING complexes, was used as an outgroup to root the tree. The support of the nodes is indicated with black dots, closed or open depending on the level of confidence as shown on the bottom left.

**Figure S3:**
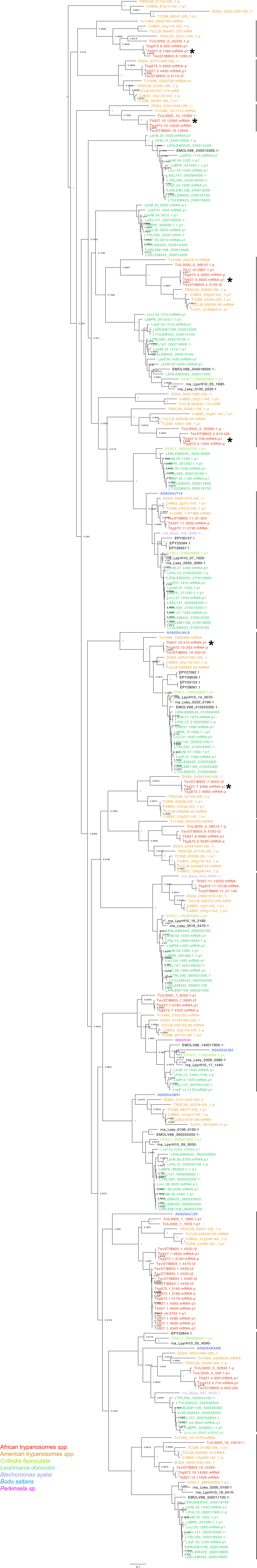
The F-box family in Kinetoplastida. Black bars mark each of the clades and stars indicate which proteins were co-precipitated with TbCul-A. Tree is rooted at midpoint and taxa is coloured coded. The Bayesian posterior probability is indicated for each node.

**Figure S4:**
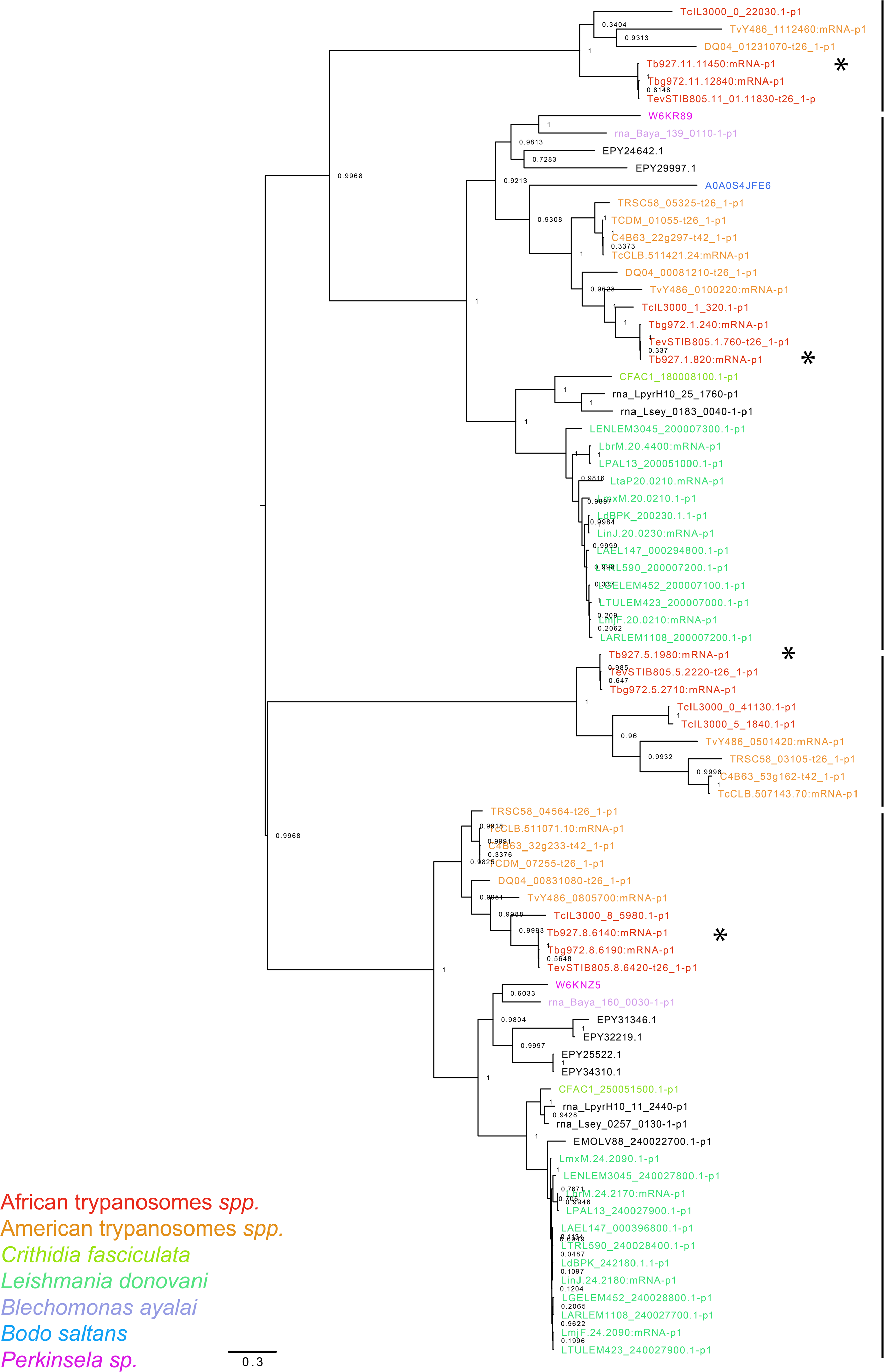
DCAF substrate adaptors in Kinetoplastida. Stars mark the two proteins identified in the immunoprecipitation of TbCul-D. Clades are indicated by black bars and the Bayesian posterior probability is indicated for each node.

**Figure S5:**
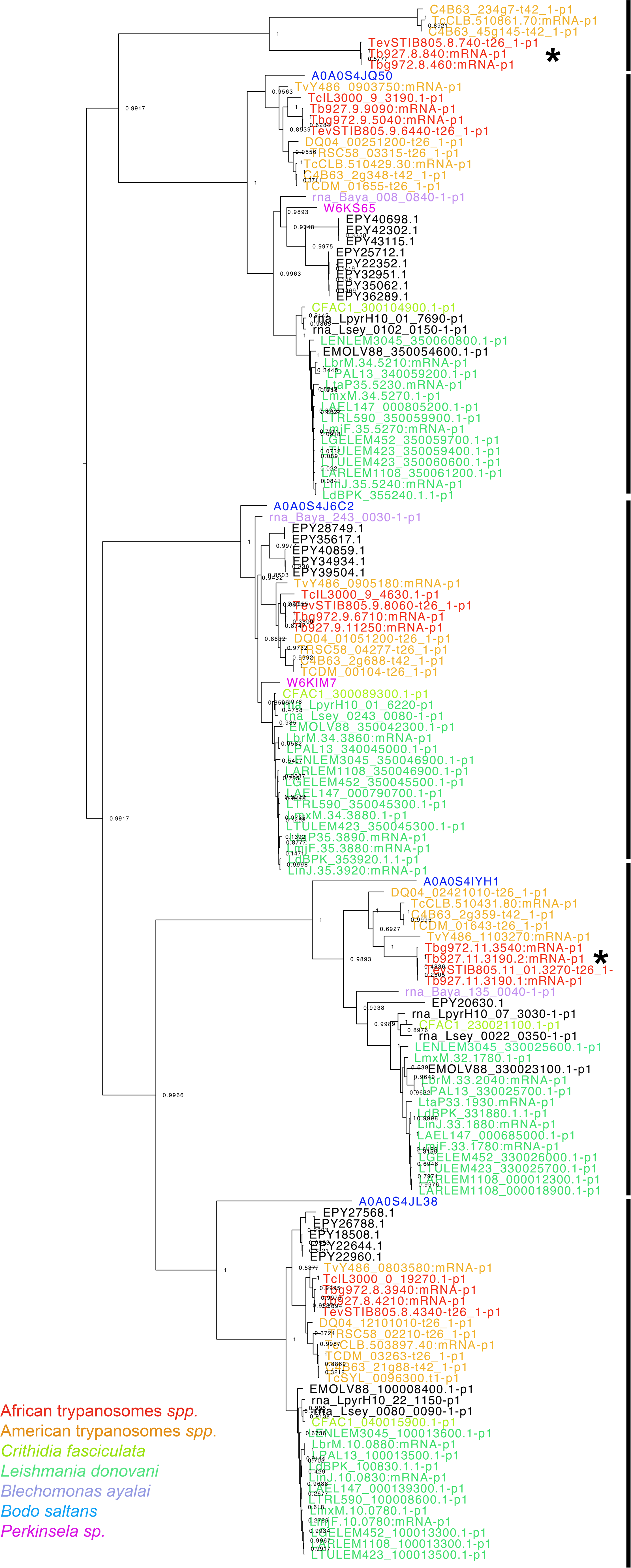
Kelch protein interactors of TbCul-E. The stars mark the four proteins identified in the immunoprecipitation of TbCul-E. Clades are indicated by black bars and the Bayesian posterior is indicated for each node.

**Figure S6:**
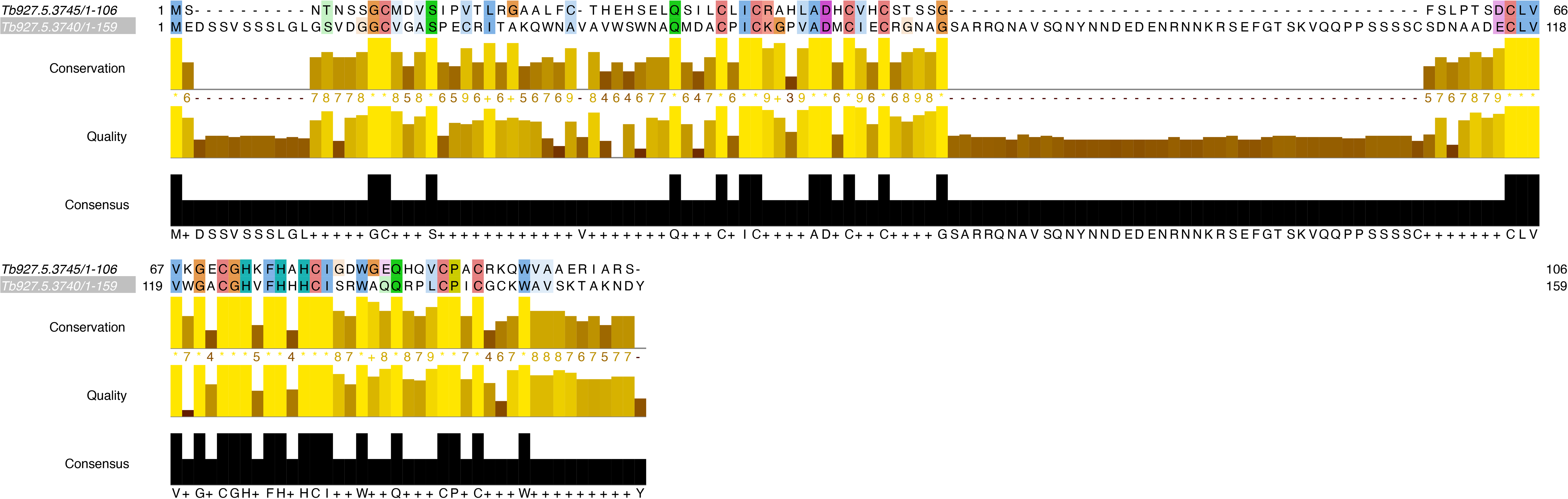
Sequence similarity of novel Rbx proteins. A cluster pairwise alignment is shown as an output from Jalview. Note that despite large indels the remaining sequences are of clear close similarity.

**Figure S7:**
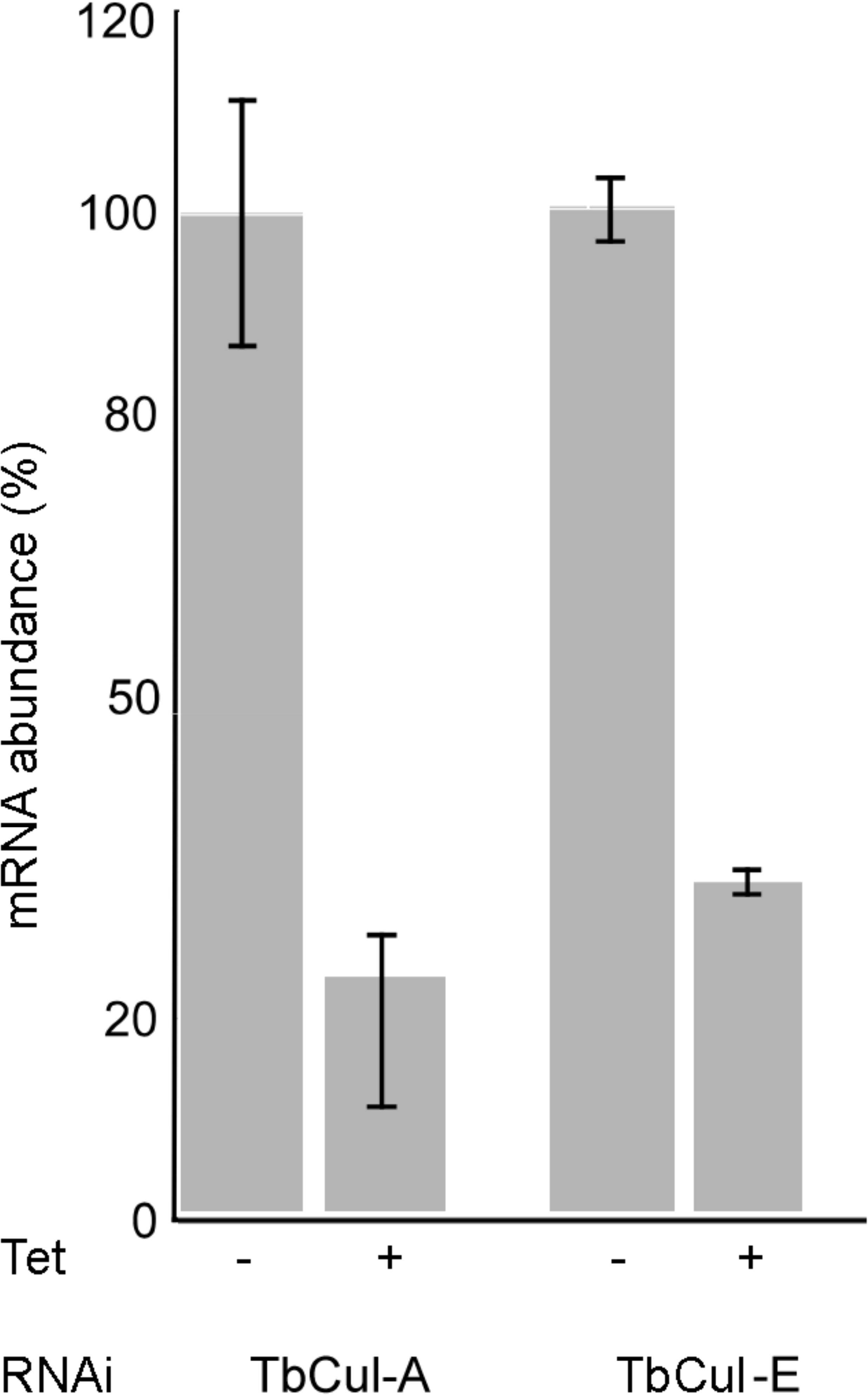
RT-qPCR validation of RNAi. RNA was extracted at the indicated times from cells induced or non induced for TbCul-A and TbCul-E knockdowns. RT-qPCR was conducted as indicated in methods. Data are from three replicate extractions for each analysis and the error bars indicate the standard deviation.

**Figure S8:**
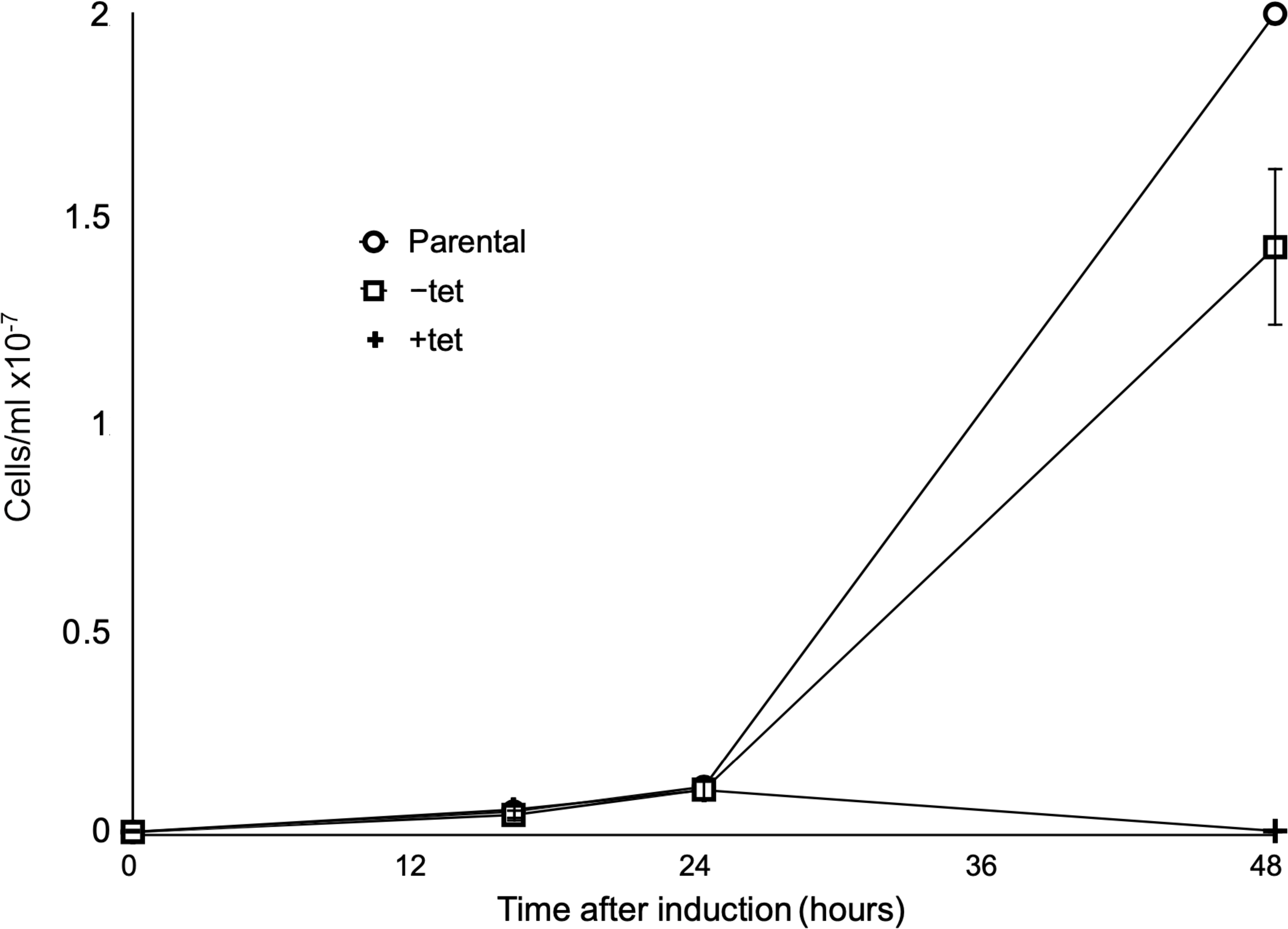
Growth curve for cells induced for TbCul-E RNAi. Cells were monitored and counted at the indicated times for three separate cultures for each condition. Error bars indicate the standard deviation.

### Supplementary tables

**Table S1:**
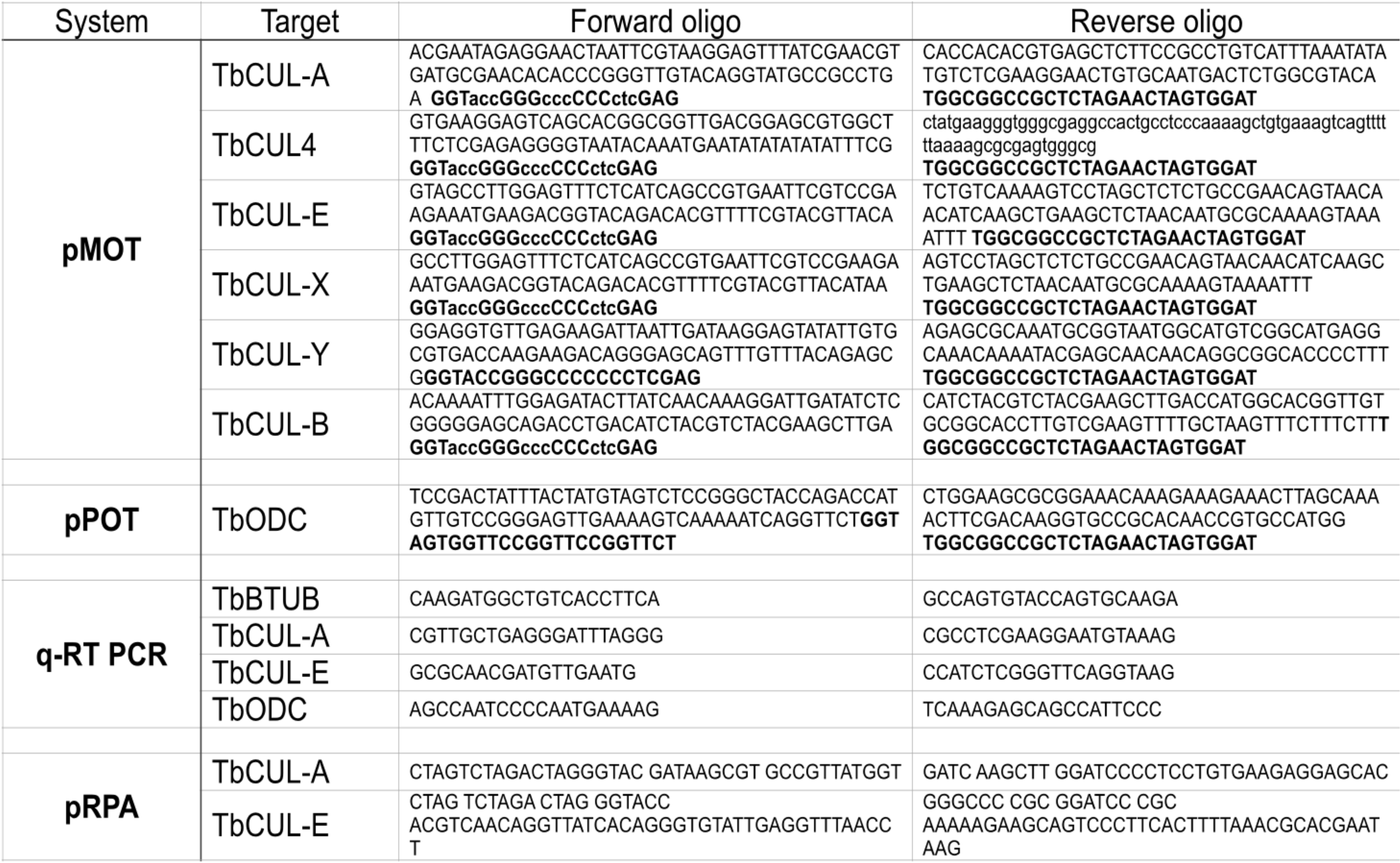
Sequences of primers used in this work. Sequences are written 5’ to 3’ and the plasmid system for which each primer was used indicated. Bold sequences are shared with pMOT and upper/lower to differentiate base triplets.

**Tables S2 – S4:** Mass spectrometry data. S2; Cullen complex immunoisolations, S3; TbCul-A knockdown, S4; TbCul-E knockdown.

**Table S5:** Species used in phylogenetic analysis.

**Supplementary data archive 1:** Fasts sequences for predicted proteins included in the phylogenetic analysis of cullins, Skp orthologs and Rbx orthologs.

